# FGF signaling regulates salivary gland branching morphogenesis by modulating cell adhesion

**DOI:** 10.1101/2022.09.10.507412

**Authors:** Ayan T. Ray, Philippe Soriano

## Abstract

Loss of FGF signaling leads to defects in salivary gland branching, but the mechanisms underlying this phenotype remain largely unknown. We disrupted expression of *Fgfr1* and *Fgfr2* in salivary gland epithelial cells and find that both receptors function coordinately in regulating branching. Strikingly, branching morphogenesis in double knockouts is restored by *Fgfr1/2* knockin alleles incapable of engaging canonical RTK signaling, suggesting that additional FGF dependent mechanisms play a role during salivary gland branching. *Fgfr1/2* conditional null mutants showed defective cell-cell and cell-matrix adhesion, both of which have been shown to play instructive roles in salivary gland branching. Loss of FGF signaling led to disordered cell-basement membrane interactions *in vivo* as well as in organ culture. This was partially restored upon introducing *Fgfr1/2* wild type or signaling alleles incapable of eliciting canonical intracellular signaling. Together, our results identify non-canonical FGF signaling mechanisms that regulate branching morphogenesis through cell adhesion processes.

## INTRODUCTION

The Sub-mandibular and the Sub-lingual Glands (SMG and SLG) regulate most of the saliva production in humans and have been widely studied in the context of branching morphogenesis (Larsen et al., 2010; Patel et al., 2006). In the mouse embryo, a stratified epithelial bud encased within a basement membrane (BM) starts to branch at embryonic (E) day 13. Progressive branching continues through E18.5 and requires instructive signals from the stromal mesenchyme to direct bud expansion followed by recurrent clefting, a feature shared with other branching organs (Costantini and Kopan, 2010; Larsen et al., 2010; Moskwa et al., 2022; Wang et al., 2017). However, *ex vivo* experiments suggest the salivary gland epithelium has an intrinsic branching potential, provided appropriate instructive signals are available (Ewald et al., 2008; Nogawa and Takahashi, 1991; Takahashi and Nogawa, 1991).

FGF signaling has been implicated in multiple steps of salivary gland branching. Our current understanding implicates FGF signaling primarily in mesenchymal-epithelial interactions (Chatzeli et al., 2017; Makarenkova et al., 2009; Sakakura et al., 1976; Steinberg et al., 2005; Wei et al., 2007). *Fgf10^-/-^* mutants exhibit branching defects in multiple exocrine organs (Hoffman et al., 2002; Jaskoll et al., 2005; Mailleux et al., 2002; Makarenkova et al., 2000; May et al., 2016; May et al., 2019; Ohuchi et al., 2000). *Fgf8* hypomorphic mutants also exhibit SMG hypoplasia (Jaskoll et al., 2004). Although SMG development remains unaffected in *Fgf7^-/-^* mutants, *ex vivo* culture experiments have shown that FGF7 can potentially regulate SMG branching (Guo et al., 1996; Steinberg et al., 2005). The FGF receptors (FGFRs) FGFR1 and FGFR2 have been thought to function in mesenchymal or epithelial contexts, respectively, however they can be co-expressed and recent reports have shown they can have combinatorial functions (Ornitz and Itoh, 2022; Ray et al., 2020). Loss of the *Fgfr2IIIb* epithelial isoform leads to acute defects in SMG development (De Moerlooze et al., 2000). More subtle effects in SMG branching have been observed by inhibiting *Fgfr1* expression *ex vivo* (Hoffman et al., 2002). In humans, haploinsufficiency of *Fgf10* and *Fgfr2* has also been implicated in autosomal dominant salivary and lacrimal gland aplasia (ALSG, OMIM 180920 and OMIM 602115 and LADD OMIM 149730) (Entesarian et al., 2007; Entesarian et al., 2005; Milunsky et al., 2006; Nie et al., 2006; Shams et al., 2007). Previous studies have shown that FGF signaling has an important function in the maintenance of epithelial progenitors in the SMG and is critical for branching (Chatzeli et al., 2017).

FGF activation engage canonical intracellular pathways, including ERK1/2, PI3K, PLCγ, JAK-STAT, JNK and p38 (Brewer et al., 2016). Inhibitor studies have implicated ERK1/2 and PI3K in SMG branching downstream of FGF pathway and other receptor tyrosine kinases (RTKs) (Kashimata et al., 2000; Larsen et al., 2003). However, FGFRs can also engage non-canonical functions (Clark and Soriano, 2022; Ray et al., 2020). Previous studies from our laboratory have revealed that *Fgfr1* and *Fgfr2* alleles lacking all canonical RTK signaling still retain extensive kinase-dependent biological activity (Brewer et al., 2015; Ray et al., 2020). FGFRs can interact through their extracellular domain with cell adhesion molecules (Doherty and Walsh, 1996; Francavilla et al., 2007; Latko et al., 2019; Nguyen and Mege, 2016; Sanchez-Heras et al., 2006; Williams et al., 1994) or heparan sulfate proteoglycans (HSPGs) which regulate BM dynamics during SMG development (Makarenkova et al., 2009). Laminin and Collagen IV act as indispensable BM components in the salivary gland (Miner and Yurchenco, 2004). *Lama5^null^* mutants develop BM defects and attenuated SMG branching (Kadoya et al., 1995; Kadoya and Yamashina, 1993; Rebustini et al., 2007). Laminin in the BM is a crucial substrate for integrin signaling and *ex vivo* culture experiments have shown that attenuation of integrin signaling function can lead to defects in salivary gland branching (Kadoya et al., 1995; Kadoya and Yamashina, 1993; Kashimata and Gresik, 1997; Wang et al., 2017). Likewise, Collagen IV has been implicated in the same process by explant culture experiments in which collagenase treatment led to defects in branching (Wang et al., 2021).

It is unknown exactly how FGFs and downstream signaling pathways regulate salivary gland epithelial branching. Previous studies using *ex vivo* culture experiments have not fully delineated FGF signaling functions in the stromal mesenchyme versus the salivary gland epithelium. In this manuscript we explore epithelial specific FGF signaling roles during salivary gland branching. Using conditional deletion of *Fgfr1* and *Fgfr2* in the salivary gland epithelial tissue, we find that both receptors function coordinately to regulate branching morphogenesis. Surprisingly, we found that these receptors regulate this process not through canonical RTK signaling, but by impinging on cell matrix interactions that regulate branching morphogenesis. Our studies highlight novel functions for FGF signaling during salivary gland development.

## RESULTS

### Spatial distribution of *Fgfr1* and *Fgfr2*

Previous research has shown that FGF signaling plays a crucial role during submandibular gland (SMG) branching. To analyze FGF receptor expression level and heterogeneity, we mined a previously published E13 mouse SMG scRNA-Seq dataset (GSE159780) (Wang et al., 2021). Based on selective marker expression followed by epithelial cell re-analysis, we identified the seven subpopulations originally visualized by the authors using graph-based dimensional Uniform Manifold Approximation and Projection (UMAP) reductions (Figure 1A). This analysis showed that among the four FGF receptors, *Fgfr1* and *Fgfr2* were predominantly expressed in the SMG epithelial cells, with both receptors showing significant heterogeneity in terms of expression levels (Figure 1A). *Fgfr3* and *Fgfr4* were expressed at significantly lower levels and were not further considered in the course of this study (Supplementary Figure 1A, B.

**Figure 1:**
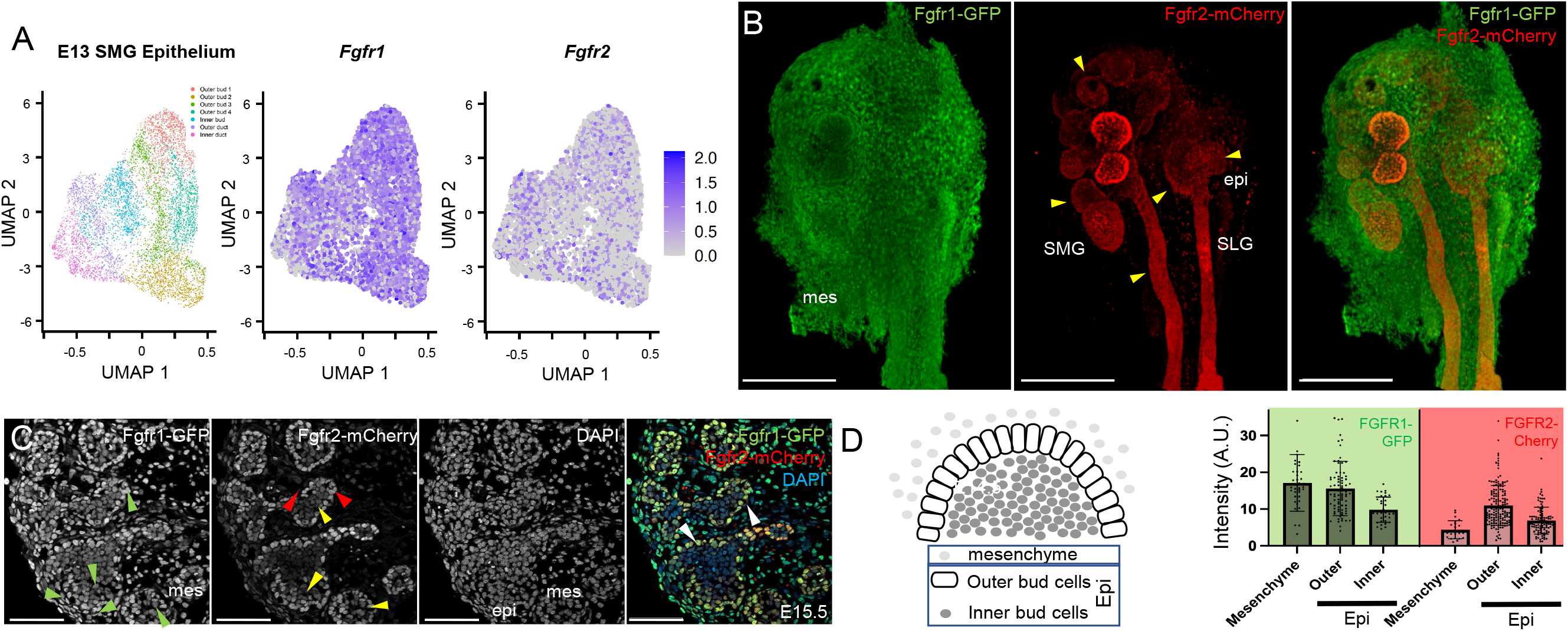
Expression of *Fgfr1* and *Fgfr2* in salivary gland. (A) Scatterplot of single cell transcriptome from E13 mouse salivary gland epithelium in the GSE159780 dataset. UMAP embedding and color coding represent 7 clusters originally identified (Wang et al., 2021). Each dot represents a single cell. Scatterplots in UMAP embedding were also color coded by expression levels for *Fgfr1* and *Fgfr2*. *Fgfr1* was more broadly expressed than *Fgfr2,* suggesting that both receptors might play important roles in SMG branching. (B) GFP and mCherry immunofluorescence shows *Fgfr1-GFP (Fgfr1)* and *Fgfr2-mCherry (Fgfr2)* expression from reporter alleles on whole mounts. Broad *Fgfr1* expression was observed in epithelial cells (epi) and the stromal mesenchyme (mes) at E13.5. *Fgfr2* expression was predominantly restricted to epithelial cells (yellow arrowheads) both in the submandibular (SMG) and sublingual glands (SLG). Scale bars, 200 □m. (C) The expression of *Fgfr1-GFP* and *Fgfr2-mCherry* was analyzed on sections at E15.5. Broad *Fgfr1* expression (green arrowheads) was observed in the mesenchyme, peripheral (outer bud) and core (inner bud) epithelial cells. Strong *Fgfr2* expression (red arrowheads) was restricted in the outer bud cells in the SMG. A low and heterogenous *Fgfr2* expression was observed in inner bud cells (yellow arrowheads). Numerous peripheral and core epithelial cells co-expressed both *Fgfr1* and *Fgfr2* (white arrowheads). Scale bars, 100 □m. (D) Schematic showing relative location of stromal/ mesenchymal cells (mesenchyme), and epithelial (Epi), outer bud (Outer) and inner bud (Inner) cells. Integrated fluorescence intensity for *Fgfr1* and *Fgfr2* were quantified for stromal (Mesenchyme), peripheral epithelial cells (Outer) and inner core epithelial cells (Inner) at E15.5 for 80-120 cells in each domain. The bar graph represents the mean integrated fluorescence (+/− SD) from *Fgfr1-GFP* and *Fgfr2-mCherry*.

We next used fluorescent *Fgfr1-GFP* and *Fgfr2-mCherry* reporter alleles (Molotkov et al., 2017) to analyze spatial domains of expression in the salivary glands. At E14.5, when several rounds of branching have occurred and the acinar program has initiated, *Fgfr1* was broadly expressed in the SMG and SLG epithelial cells as well as in mesenchyme/stroma. In contrast, *Fgfr2* expression was restricted to epithelial cells (Figure 1B). The outer bud epithelial cells, which make strong cell-basement membrane interactions and play important roles in remodeling the BM during branching, expressed higher levels of both *Fgfr1* and *Fgfr2.* Outer bud cells subsequently lose cell-basement membrane contacts during epithelial expansion and populate the inner core (Wang et al, 2021). We further analyzed expression of *Fgfr1-GFP* and *Fgfr2-mCherry* on sections at E15.5. Broad *Fgfr1* expression was observed at this stage in both outer and inner bud epithelial and stromal (mesenchymal) cells. Strong *Fgfr2* expression was found primarily in the outer bud epithelial cell population, similar to *Fgfr1*. The majority of inner bud cells co-expressed *Fgfr1* and *Fgfr2* at generally lower and heterogeneous levels (Figure 1C, D). Together, our analysis showed that both FGF receptors are dynamically expressed during SMG development, with high levels of both receptors in outer bud epithelial cells while heterogenous expression was observed in inner core epithelial cells.

### *Fgfr1* and *Fgfr2* regulate SMG branching

Salivary glands progressively branch between E13 and E18.5. To further understand the role of *Fgfr1* and *Fgfr2* in epithelial cells during branching morphogenesis, we combined a *K14Cre* driver active in the SMG (Lombaert et al., 2013) with conditional null alleles for *Fgfr1* and *Fgfr2* (henceforth referred to as *cKO*) to attenuate FGF signaling in the salivary gland epithelium and ductal progenitors (Supplementary Figure 2A, B). Tissues were harvested at E14.5, followed by immunostaining of epithelial cells for E-cadherin (ECAD) in whole mounts, and the number of terminal buds were counted for conditional mutants of *Fgfr1* (Figure 2A and, D), *Fgfr2* (Figure 2B, D) and compound *Fgfr1/2* double mutants (Figure 2C, D), as well as Cre negative controls from ECAD 3D rendered images. *Fgfr1^+/cKO^* mutants did not develop branching defects. However, a 43% reduction in the number of terminal buds was observed in *Fgfr1^cKO/cKO^* mutants (Figure 2A, D). The SLG remained unaffected in these mutants. Interestingly, *Fgfr2^+/cKO^* mutants developed a slight reduction in terminal bud numbers at E14.5. *Fgfr2^cKO/cKO^* mutants exhibited a large reduction in the number of SMG terminal buds and also did not develop a SLG (Figure 2). This suggests the two receptors may regulate specific aspects of branching. Interestingly, the conditional *Fgfr2^cKO/cKO^* mutant phenotype was less severe than *Fgfr2IIIb^null^* mutants, which survive through mid-gestation but do not develop any salivary glands and die at birth (De Moerlooze et al., 2000). Since all epithelial lineages in the salivary gland originate from *Krt14^+^* lineages (May et al., 2018), *Fgfr2* might have earlier functions in the oral ectoderm or in progenitor maintenance. Lastly, we observed that *Fgfr1^+/cKO^; Fgfr2^cKO/cKO^* double conditional mutants showed the most severe phenotype with only one to four terminal buds (Figure 2C, D), indicating that both receptors have combinatorial functions during epithelial branching. The fact that both receptors are co-expressed throughout the SMG cells (Figure 1C, D) also suggests that they might have overlapping roles. The presence of a single wild-type *Fgfr1* or *Fgfr2* allele could partially rescue branching defects in *Fgfr1^cKO/cKO^; Fgfr2^cKO/cKO^* compound conditional mutants. *Fgfr1^+/cKO^; Fgfr2^cKO/cKO^* compound conditional mutants rescued terminal branching defects by 19.2%. A more robust rescue (~71%) in terminal branching was observed in *Fgfr1^cKO/cKO^; Fgfr2^+/cKO^* mutants, again supporting a more extensive role for *Fgfr2* in branching morphogenesis.

**Figure 2:**
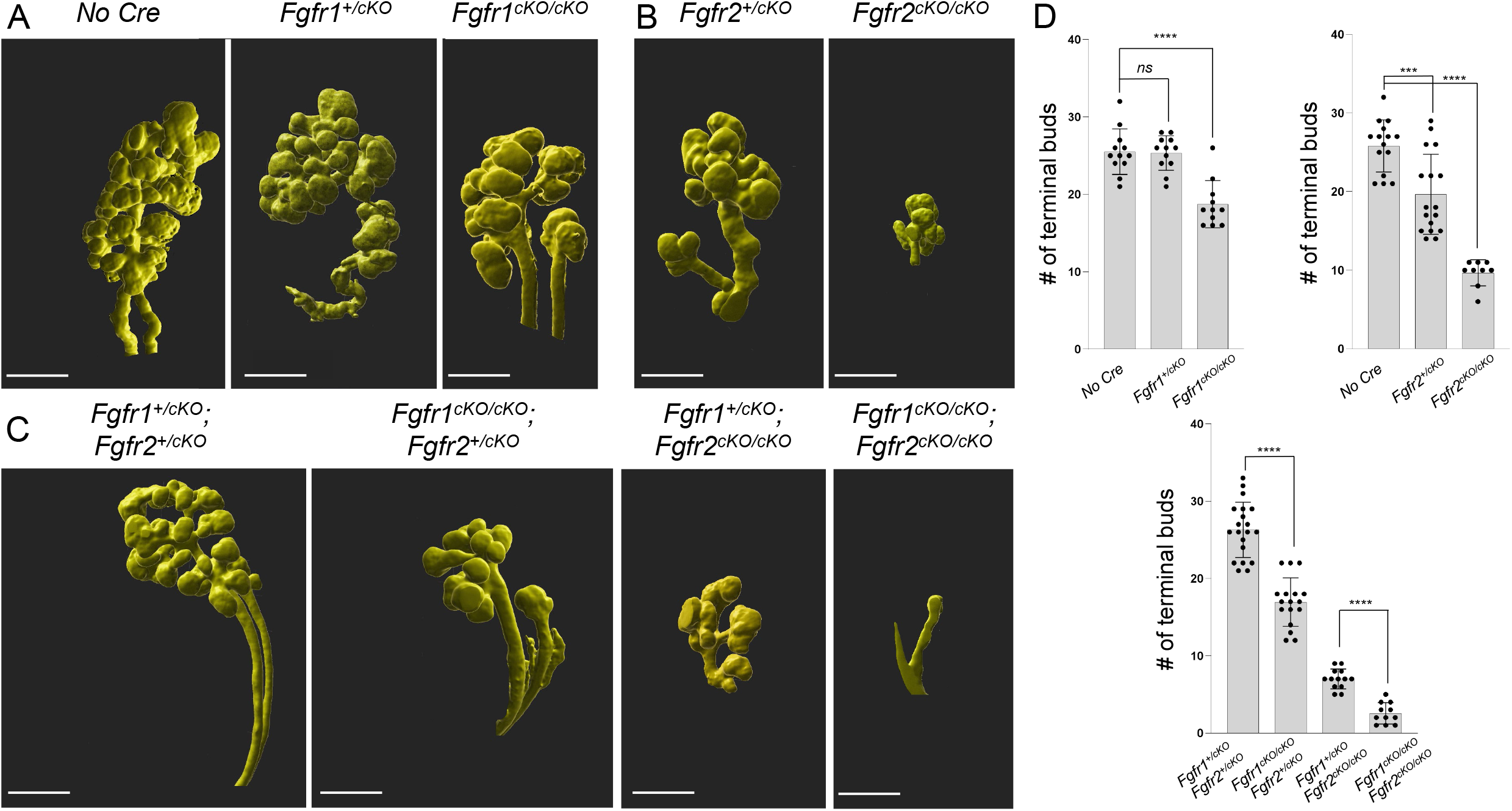
Branching defects in *Fgfr1* and *Fgfr2* conditional mutants. Conditional depletion of (A) *Fgfr1,* (B) *Fgfr2,* or (C) compound *Fgfr1/2* using *K14Cre* resulted in branching defects at E14.5 in developing salivary glands. Wholemount E-cadherin immunofluorescence followed by 3D rendering of high-resolution images was used to visualize these defects. The number of terminal buds across various *Fgfr1* and *Fgfr2* conditional mutants were analyzed. Severe defects in SMG branching were observed in compound *Fgfr1^cKO/cKO^; Fgfr2^cKO/cKO^* mutants where number of buds varied between 1 and 5. Conditional *Fgfr2^cKO/cKO^* mutants, *Fgfr1^+/cKO^; Fgfr2^cKO/cKO^* mutants and *Fgfr1^cKO/cKO^; Fgfr2^cKO/cKO^* mutants did not develop an SLG. Scale bars, 200 μm. (D) Bar graphs show the number of terminal buds in the SMG for various conditional *Fgfr1* and *Fgfr2* mutants.

### Canonical FGF signaling does not affect SMG branching

We sought to understand if branching defects in various *Fgfr1/2* compound mutants result from changes in cell proliferation or survival, two processes known to be regulated by FGF signaling. We assessed cell proliferation by EdU incorporation across various *Fgfr1/2* conditional mutants at E14.5 when defects in branching already appeared. Control or compound *Fgfr1/2* mutants showed similar levels of EdU incorporation in both the epithelium as well as the stromal mesenchyme (Supplementary Figure 3A). Likewise, we found little cell death and no increase in single or compound conditional *Fgfr1/2^cKO/cKO^* mutants at this stage by TUNEL staining (Supplementary Figure 3B). A defect in proliferation or cell death may be present at earlier stages, however, as *Fgfr1^+/cKO^; Fgfr2^+/cKO^* controls showed many more K14 lineage GFP^+^ cells than mutants. Alternatively, it is possible that loss of FGF receptors affects the recruitment of epithelial progenitors from the oral ectoderm to the salivary glands.

FGF receptors can engage multiple signaling pathways, including ERK1/2, PI3K/Akt, PLCγ and STATs (Brewer et al., 2016). Previous pharmacological inhibitors studies in *ex vivo* explant cultures have implicated ERK1/2 activation during epithelial branching (Kashimata et al., 2000). To further explore the contribution of ERK1/2 activation downstream of FGF receptors, we used a conditional constitutively-active *Braf* allele *(BrafCA)* allele to induce the RAF-MEK-ERK pathway in the *K14Cre* lineage independent of *Fgfr1/2* expression. We observed significant defects in branching in *BrafCA* mutants in a *Fgfr1^cKO/cKO^; Fgfr2^cKO/cKO^ null* background. RAF-MEK-ERK activation in *Fgfr1^cKO/cKO^; Fgfr2^cKO/cKO^; BrafCA* mutants (Supplementary Figure 3C) could not alleviate branching defects observed upon elimination of *Fgfr1* and *Fgfr2* (Figure 3A). These observations suggest that that ERK1/2 activation is not sufficient to rescue defects caused by loss of FGF signaling.

**Figure 3:**
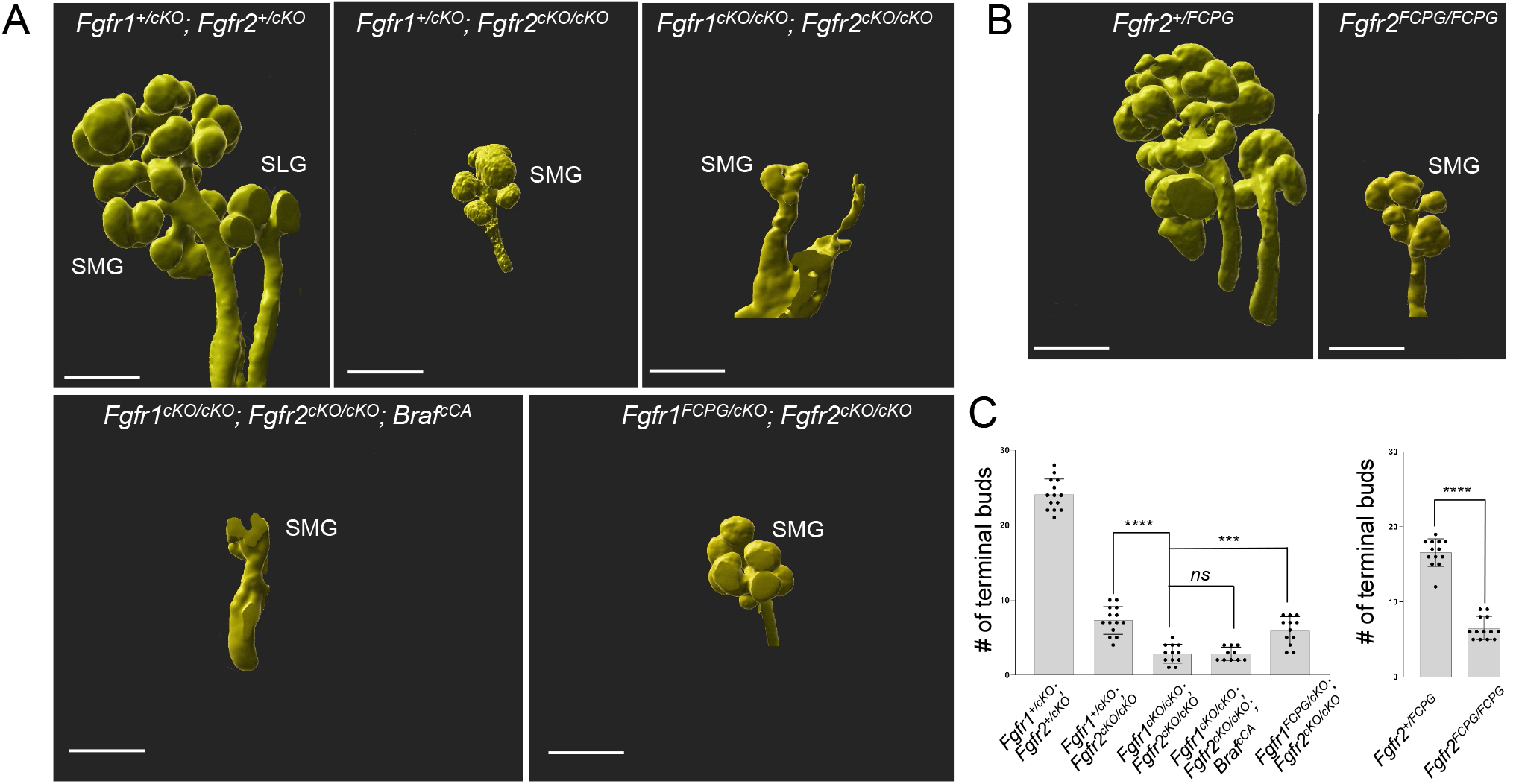
Branching in signaling mutants. (A) Wholemount E-cadherin immunofluorescence followed by 3D rendering was used to analyze changes in number of terminal buds at E14.5 across various *Fgfr1/2* signaling mutants. *Fgfr1^cKO/cKO^; Fgfr2^cKO/cKO^; BrafCA* mutants did not rescue branching defects in the SMG. The SLG did not develop in these mutants. Compared to severe branching defects observed in *Fgfr1^cKO/cKO^; Fgfr2^cKO/cKO^* mutants, *Fgfr1^FCPG/cKO^; Fgfr2^cKO/cKO^* mutants exhibited a partial rescue of branching defects in the SMG. SLG did not develop in *Fgfr1^FCPG/cKO^; Fgfr2^cKO/cKO^* mutants. A representative image is shown for each genotype. Scale bars, 200 μm. (B) 3D rendering was used to analyze SMG defects in *Fgfr2^+/FCPG^* and *Fgfr2^FCPG/FCPG^* mutants at E14.5. *Fgfr2^FCPG/FCPG^* mutants did not develop sublingual glands and showed reduced terminal bud numbers in the SMG. Scale bars, 200 μm. (C) The mean terminal bud number (+/− SD) in the SMG for various *Fgfr1* and *Fgfr2* signaling mutants at E14.5 were represented as bar graphs. *Fgfr1^cKO/cKO^; Fgfr2^cKO/cKO^* mutants showed a 94.5% reduction in terminal bud numbers. A similar level of reduction was observed upon ectopic activation of BRAF in *Fgfr1^cKO/cKO^; Fgfr2^cKO/cKO^; Braf^cCA^* mutants. *Fgfr1^FCPG/cKO^; Fgfr2^cKO/cKO^ mutants* rescued branching defects by 19.1%. A 61% reduction in terminal bud number was observed in *Fgfr2^FCPG/FCPG^* mutants compared to controls.

Since ERK1/2 activation did not rescue epithelial branching defects in *Fgfr1/2* compound mutants, we wanted to explore if other FGF signaling functions mediate this process. To interrogate signaling mechanisms downstream of FGF signaling *in vivo,* we previously generated mice with knock-in point mutations at the *Fgfr1* and *Fgfr2* loci that prevent binding of effectors to the receptors. The most severe alleles, *Fgfr1^FCPG^* and *Fgfr2^FCPG^*, removed all canonical intracellular signaling downstream of the receptors, including ERK1/2, PI3K, PLC g and STAT signaling, but failed to recapitulate null mutant phenotypes (Brewer et al., 2016; Brewer et al., 2015; Ray et al., 2020). To investigate the role of canonical FGF intracellular signaling activation during branching, we analyzed *Fgfr1^FCPG/cKO^; Fgfr2^cKO/cKO^* and *Fgfr2^FCPG/FCPG^* mutants.

Whole-mount ECAD immunostaining followed by 3D rendering of images was used to determine changes in terminal number of buds in compound signaling mutants at E14.5. *Fgfr1^cKO/cKO^; Fgfr2^cKO/cKO^* mutants showed pronounced branching defects as described earlier. *Fgfr1^FCPG/cKO^; Fgfr2^cKO/cKO^* mutants showed a partial rescue (19.1% compared to *Fgfr1^cKO/cKO^; Fgfr2^cKO/cKO^* compound conditional mutants) in terminal bud formation, to nearly the same extent as a wild type *Fgfr1* allele. *Fgfr1^FCPG^* allele did not affect defects observed in SLG development. Morphologically, branching defects in *Fgfr1^FCPG/cKO^; Fgfr2^cKO/cKO^* compound conditional mutants phenocopied *Fgfr1^+/cKO^; Fgfr2^cKO/cKO^* mutants (Figure 3A, C). Thus, both *Fgfr1^FCPG/cKO^; Fgfr2^cKO/cKO^* and *Fgfr1^+/cKO^; Fgfr2^cKO/cKO^* compound conditional mutants, showed similar rescue of branching defects as determined by terminal end bud number upon loss of FGFR1/2 functions.

Likewise, *Fgfr2^FCPG/FCPG^* mutants exhibited partial rescue of terminal bud formation in contrast to *Fgfr2III^b^* mutants that show agenesis of the salivary glands (Figure 3B, C)(Celli et al., 1998; De Moerlooze et al., 2000). Interestingly, defects in SLG formation were still not rescued in *Fgfr2^FCPG/FCPG^* mutants (Figure 3B). We found that the SLG never formed in *Fgfr2^FCPG/FCPG^* adult mice, but that the SMG was similarly sized as controls suggesting that branching morphogenesis catches up after E14.5. We were unable to analyze conditional mutants in combination with the *Fgfr2^FCPG^* allele due to *loxP* sites carried over from the sequential introduction of the FCPG mutations. Taken together, these results indicate that both *Fgfr1^FCPG^* and *Fgfr2^FCPG^* alleles can significantly rescue *Fgfr1^-/-^ and Fgfr2^-/-^* mutant branching morphogenesis defects and that both receptors must function in part through other mechanisms than canonical signaling in salivary glands.

### BM defects in *Fgfr1/2* mutants

The BM surrounding the salivary gland has been shown to play critical roles instructing epithelial remodeling and SMG branching, in part governed by the strength of outer bud cellmatrix adhesion (Wang et al., 2021). Since FGFRs have functions in regulating cell adhesion beyond their classical roles in canonical signaling (Ray et al., 2020), we turned our attention to interactions between outer bud cells and the BM, which are critical for SMG branching morphogenesis. We investigated the dynamics of epithelial branching in the SMG between E13.5 and E15.5 when epithelial buds are most significantly remodeled. Branching has been shown to occur by proliferation of the terminal buds and recurrent clefting events during outgrowth. We carried out a time course analysis using ECAD whole mount immunostaining to label the epithelial sheet and measure the average curvature as development progresses. This analysis provided a quantitative perspective regarding the extent of epithelial remodeling (Figure 4A, A’).

**Figure 4:**
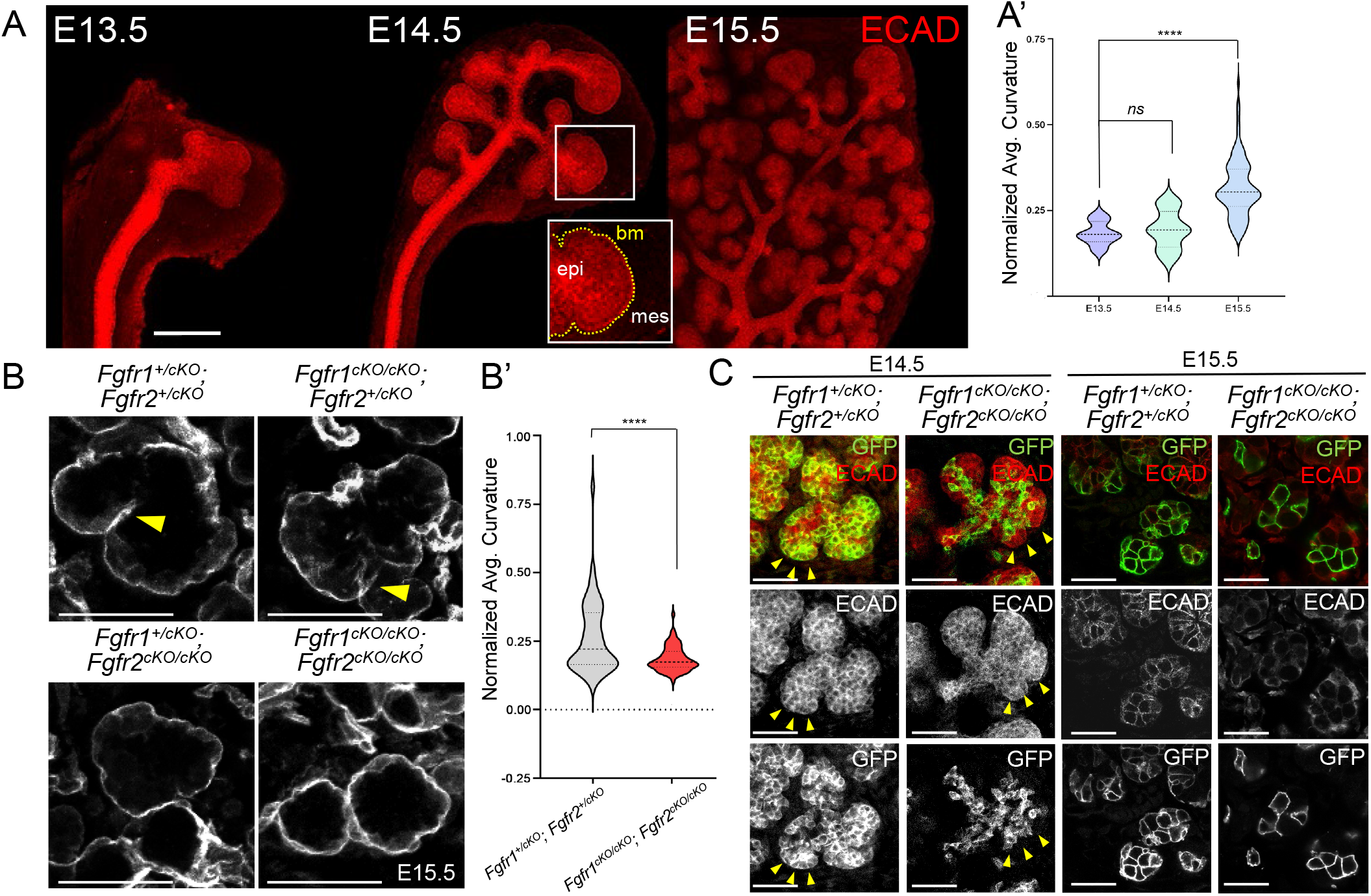
Defects in BM in *Fgfr1/2* mutants. (A) E-cadherin immunofluorescence was used to analyze SMG epithelial branching between E13.5 to E15.5 in control SMGs. The epithelial surface (epi) sheet is enclosed by a basement membrane (bm) that undergoes dramatic folding within the mesenchyme (mes), due to recurrent clefting and outgrowth. The median of average curvature of the epithelial surface is represented as violin plots (A’) from E13.5 to E15.5. The horizontal bold dashed bar represents the median. The lighter bars represent the first and third quartiles. Scale bar 200 μm. (B) BM of SMG was labeled by Laminin immunofluorescence on sections for *Fgfr1/2* compound mutants at E15.5 to estimate the extent of branching. Numerous invaginations (yellow arrowheads) that represent independent clefting events, were detected in control *Fgfr1^+/cKO^* and *Fgfr2^+/cKO^* and *Fgfr1^cKO/cKO^; Fgfr2^+/cKO^* mutant SMGs. A significant reduction in clefting was observed in *Fgfr1^+/cKO^; Fgfr2^cKO/cKO^* mutants and *Fgfr1^cKO/cKO^; Fgfr2^cKO/cKO^* compound mutant SMGs. Median average curvature for Laminin surface is shown as violin plots for controls and *Fgfr1^cKO/cKO^; Fgfr2^cKO/cKO^* mutants (B’). Scale bars, 50 μm. (C) Compound *Fgfr1/2* mutant SMG were analyzed at E14.5 and E15.5. GFP^+^*Fgfr1^cKO/cKO^; Fgfr2^cKO/cKO^ K14* lineage cells were restricted to the inner ECAD^+^ epithelial domain compared to controls at E14.5. Very few GFP^+^ cells could be detected in the peripheral border region (yellow arrowheads) in *Fgfr1^cKO/cKO^; Fgfr2^cKO/cKO^* mutants at E14.5. Scale bars, 50 μm.

During branching, as the outer bud cells invaginate as a surface sheet to facilitate clefting, the BM is remodeled and the average curvature of the BM increases. We analyzed changes in BM curvature in compound *Fgfr1/2* mutants on a *ROSA26^mTmG^* background by immunostaining for Laminin on sections at E15.5. We observed that control compound *Fgfr1^+/cKO^; Fgfr2^+/cKO^* mutants developed extensive invaginations by E15.5 (Figure 4B, yellow arrowhead). *Fgfr1^cKO/cKO^; Fgfr2^+/cKO^* mutants also developed profound invaginations, however, they were more subtle in nature compared to controls. *Fgfr1^+/cKO^; Fgfr2^cKO/cKO^* mutants showed fewer clefting events along with reduced changes in BM curvature and smoother margins, (Figure 4B). The most severe defects in BM curvatures were observed in *Fgfr1^cKO/cKO^; Fgfr2^cKO/cKO^* compound mutants, as indicated by the smooth BM, and rarely showed sporadic clefting events compared to *Fgfr1^+/cKO^; Fgfr2^+/cKO^* compound heterozygotes. This is consistent with the significant reduction in average basement membrane curvature observed in compound *Fgfr1^cKO/cKO^; Fgfr2^cKO/cKO^* null mutants (Figure 4B, B’). *Fgfr1^cKO/cKO^; Fgfr2^cKO/cKO^* compound mutants consistently showed defects in cell BM interactions. At E14.5, GFP^+^*Fgfr1^cKO/cKO^; Fgfr2^cKO/cKO^* mutant cells showed a striking failure to populate the outer margins of the epithelial buds, which were instead composed of ECAD^+^; GFP^-^ cells abutting the BM, and these defects were exacerbated by E15.5 (Figure 4C).

Budding morphogenesis has been shown to be driven by strong cell-matrix adhesion, but also concomitant weak cell-cell adhesion at the periphery and strong cell-cell adhesion of inner epithelial cells (Wang et al., 2021). We observed that while *K14Cre* labeled GFP^+^ cells populated salivary gland acini extensively and exhibited strong cell-cell adhesion marked by ECAD staining in control *Fgfr1^+/cKO^; Fgfr2^+/cKO^,* far fewer cells were seen in *Fgfr1^cKO/cKO^; Fgfr2c^KO/cKO^* acini. Remarkably, these cells showed reduced ECAD accumulation at the cell-cell junctions suggesting weak cell-cell adhesion (Supplementary Figure 4A, B). Together, these results indicate that loss of FGF signaling diminishes cell-cell adhesion within the SMG. Interestingly, *Fgfr1^FCPG/cKO^; Fgfr2^cKO/cKO^* mutants maintain robust ECAD accumulation at the cell-cell junctions comparable to *Fgfr1^+/cKO^; Fgfr2^+/cKO^* controls (Supplementary Figure 4C), indicating that cell-adhesion regulation is controlled by non-canonical FGF signaling.

The basement membrane that surrounds the epithelial end buds becomes remodeled during branching morphogenesis and is perforated at areas of local expansion to allow further growth (Harunaga et al., 2014). We examined BM integrity in *Fgfr1/2* conditional *null* mutants on sections. In control *Fgfr1^+/cKO^; Fgfr2^+/cKO^* mutants, the basal edges of GFP^+^ cells made extensive contacts with the BM. These interactions were largely compromised in *Fgfr1^cKO/cKO^; Fgfr2^cKO/cKO^* mutants (Figure 5A). *Fgfr1^cKO/cKO^; Fgfr2^cKO/cKO^* mutants also showed major disintegration and discontinuity of the BM specifically around the developing acini (Figure 5B, yellow arrowheads). We further analyzed cell BM interactions in *Fgfr1^cKO/cKO^; Fgfr2^cKO/cKO^* mutants using high resolution confocal microscopy of 50μm thick sections stained for the BM protein Laminin. Controls showed extensive co-localization of Laminin and membrane-GFP, however, this was diminished in *Fgfr1^cKO/cKO^; Fgfr2^cKO/cKO^* mutants (Supplementary Figure 5A). Laminin staining showed large areas of discontinuity in *Fgfr1^cKO/cKO^; Fgfr2^cKO/cKO^* mutant tissues, but not controls (Supplementary Figure 5B, white arrowheads). Therefore, a qualitative examination revealed that in *Fgfr1^cKO/cKO^; Fgfr2^cKO/cKO^* mutants, the BM appeared disintegrated in certain domain, with large gaps, variable thickness, and minimal interaction with GFP^+^ cells. These results indicate that *Fgfr1/2* plays important role in cell-BM interaction and in maintaining BM integrity.

**Figure 5:**
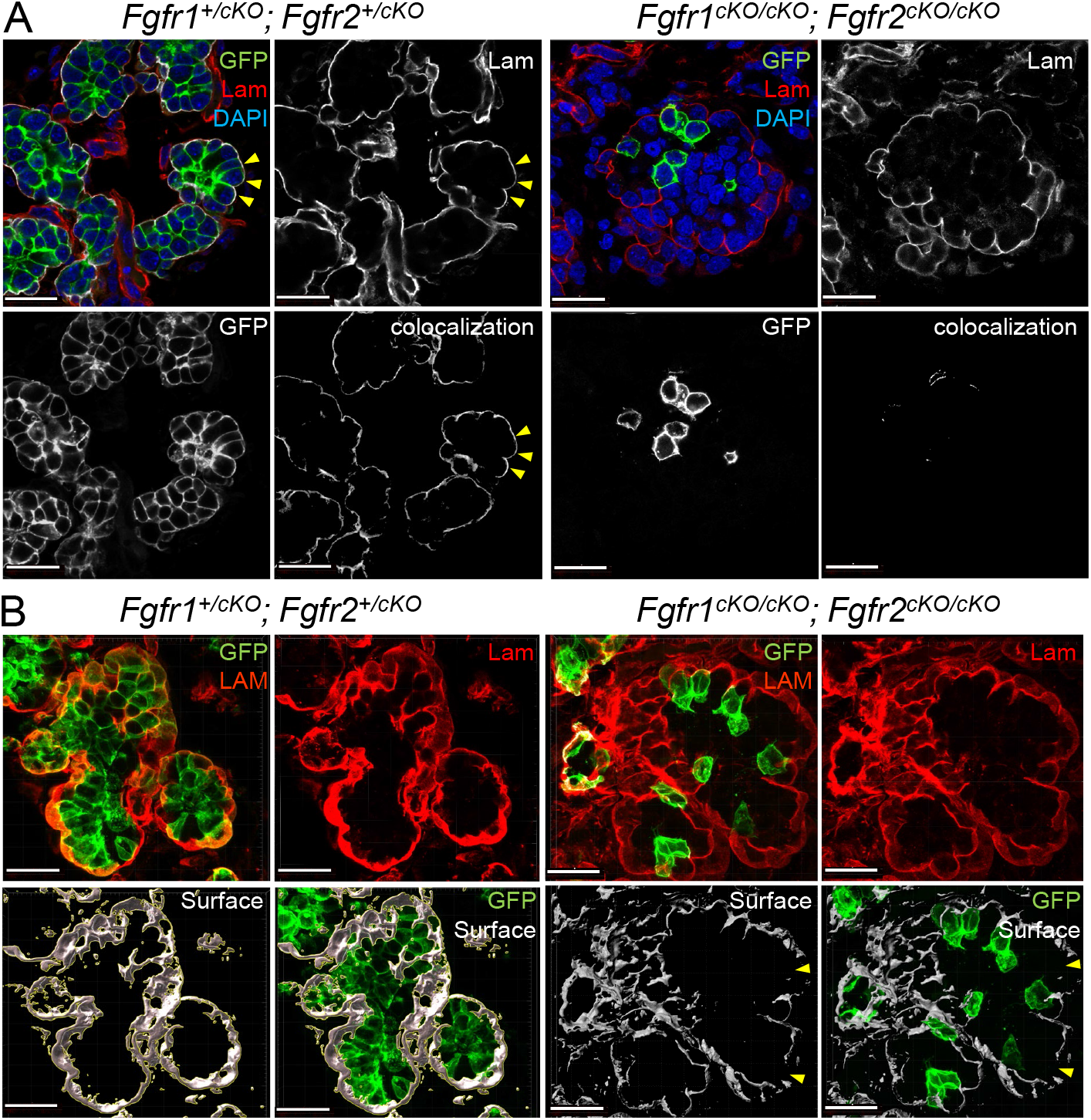
Cell-BM interactions in *Fgfr1/2* compound mutants. (A) Cell-BM interactions in the SMG were analyzed for compound *Fgfr1/2* compound mutants at E15.5 by generating 3D colocalization maps. The BM was marked by Laminin (Lam). *K14* lineage epithelial cells expressed GFP. High resolution confocal imaging followed by 3D rendering was used to detect colocalization between GFP of Lam and colocalization maps were generated. Control cells formed robust cell-BM contacts (yellow arrowheads). Several GFP^+^, *Fgfr1^cKO/cKO^; Fgfr2^cKO/cKO^* cells did not interact with the BM. Scale bars, 20 μm. (B) Integrity of basement membrane in compound *Fgfr1/2* mutants were analyzed. 3D surfaces around Laminin were generated from high resolution 3D rendered confocal images. Compound *Fgfr1^cKO/cKO^; Fgfr2^cKO/cKO^* mutants showed discontinuity along the laminin surfaces (yellow arrowheads). Scale bars, 20 μm.

### FGF-dependent integrin functions during branching morphogenesis

Our analysis suggests that altered SMG branching in compound *Fgfr1/2* mutants might arise from defects in cell-BM interactions. The integrins ITGA6 and ITGB1 have been shown to interact with Laminin during SMG remodeling. In organ culture, anti-ITGA6 antibodies inhibit SMG branching (Kadoya et al., 1995; Kadoya and Yamashina, 1993). We further mined the published sc-RNA seq data for E13 mouse SMG (GSE159780, (Wang et al., 2021)), and found *Itga6, Itga9* and *Itgb1* to be strongly expressed in the outer bud epithelial cells (Supplementary Figure 6A). We used immunofluorescence to detect ITGB1 in wild-type salivary glands at E14.5 and E15.5 (Supplementary Figure 6B). Since integrins positively regulate branching, we wanted to investigate if ITGB1 can cooperate with or attenuate FGF signaling function during branching.

To investigate if integrin signaling activation could rescue branching defects in *Fgfr1/2* conditional mutants, we collected *Fgfr1/2* mutant SMGs at E14.5 and cultured them in media supplemented with Mn^2+^, which is known to lock ITGB1 in a ligand bound active conformation and ectopically activate integrin signaling (Bazzoni et al., 1995). We observed robust integrin signaling activation upon Mn^2+^ treatment within 8 hours of culture (Supplementary Figure 6C). Changes in branching were analyzed at 18 hrs and 40 hrs (Figure 6A, Supplementary Figure 6C). Compared to untreated culture conditions, we found that integrin activation resulted in a robust increase in the curvature index and bud number in *Fgfr1^+/cKO^; Fgfr2^+/cKO^* controls at 40 hrs. *Fgfr1^cKO/cKO^; Fgfr2^cKO/cKO^* mutants however did not show a significant increase in branching, suggesting that activation of integrin signaling is not sufficient to induce ectopic branching upon loss of both FGF receptors.

**Figure 6:**
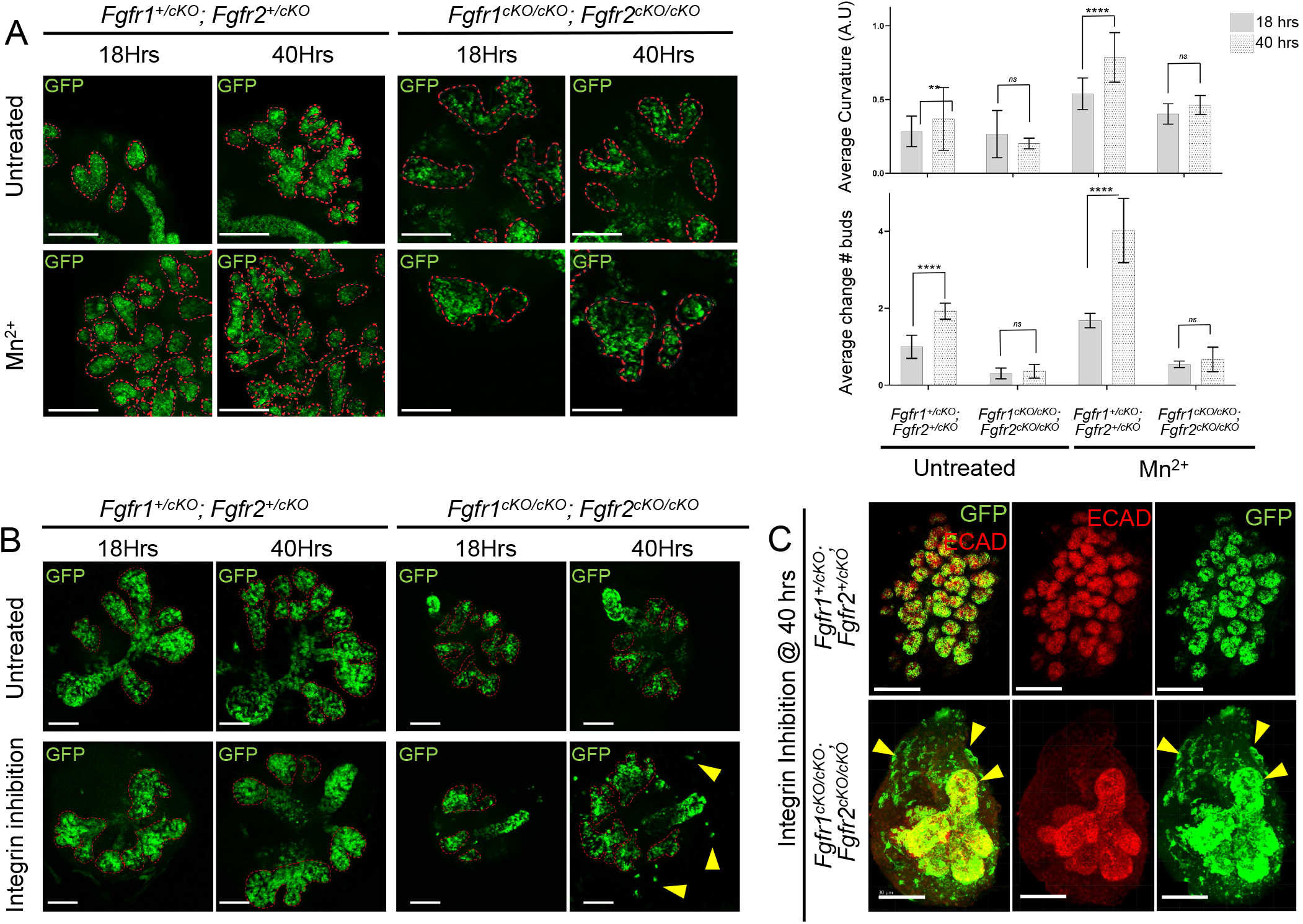
Integrin signaling cooperates with FGFR function during SMG branching. (A) SMG explants from various *Fgfr1* and *Fgfr2* compound mutants (E14.5) were cultured for 40 hrs in presence of Mn^2+^ (integrin signaling activation) and compared to untreated controls. Explants were imaged during branching at 18 hrs and 40 hrs of culture. The curvature of the epithelial surface is depicted by dashed red lines. Median average curvatures (error bar represent 95% confidence interval, CI) and average change in bud numbers (error bar represent standard deviation) were measured for each genotype and represented as a bar graph. Robust branching was observed upon integrin signaling activation in all mutants except for *Fgfr1^cKO/cKO^; Fgfr2^cKO/cKO^.* Scale bars, 100 μm. (B) SMG explants (E14.5) were treated with an integrinβ1 blocking antibody from various *Fgfr1* and *Fgfr2* compound mutants. Epithelial branching was assessed at 18 hrs and 40 hrs during culture. Median average curvature was plotted as bar graphs (Supplementary Figure S6). Neither control *Fgfr1^+/cKO^; Fgfr2^+/cKO^* nor *Fgfr1^cKO/cKO^; Fgfr2^cKO/cKO^* compound mutant explants showed robust changes in branching upon integrin signaling inhibition. However, several GFP^+^ double mutant cells extruded from the explants (yellow arrowheads). Scale bars, 100 μm. (C) Upon integrin signaling inhibition for 40 hrs, explants were immunostained for ECAD to mark epithelial cells. Compared to *Fgfr1^+/cKO^; Fgfr2^+/cKO^* controls, several GFP^+^ cells in *Fgfr1^cKO/cKO^; Fgfr2^cKO/cKO^* explants extruded out of the ECAD^+^ epithelial domain into the surrounding mesenchyme. Scale bars, 200 μm.

Next, we used a blocking antibody to inhibit integrin signaling in culture. We cultured control *Fgfr1^+/cKO^; Fgfr2^+/cKO^* and *Fgfr1^cKO/cKO^; Fgfr2^cKO/cKO^* salivary glands in the presence and absence of a blocking antibody and analyzed branching at 18 hrs and 40 hrs. Branching remained unaffected in both controls and mutants upon integrin signaling inhibition (Figure 6B, Supplementary Figure 6E). However, we observed extensive extrusion of GFP^+^ cells into the surrounding mesenchyme in *Fgfr1^cKO/cKO^; Fgfr2^cKO/cKO^* mutants at the 40 hrs endpoint, exclusively in the presence of the integrin blocking antibody (Figure 6B, C, yellow arrowhead). We further analyzed these mutants on sections. In control *Fgfr1^+/cKO^; Fgfr2^+/cKO^* explants, we observed extensive remodeling of the BM in both the untreated and integrin-blocking conditions, and GFP^+^cells made extensive contacts with the BM in both conditions. *Fgfr1^cKO/cKO^; Fgfr2^cKO/cKO^* compound mutants did not show overt defects in the BM in the untreated condition, but upon integrin inhibition the BM showed drastically reduced Laminin levels and GFP^+^ cells often had an abnormal shape, appearing more elongated than round or cuboidal as in the untreated condition (Supplementary Figure 6E). These results suggest that integrin inhibition further exacerbates *Fgfr1/2* mutant defects.

### FGF receptor signaling mutants rescue branching defects upon integrin function activation

We earlier observed that integrin signaling activation could not rescue branching defects in *Fgfr1/2* compound *null* mutants (Figure 6A). To determine if canonical *Fgfr1* and *Fgfr2* signaling plays a role during branching upon integrin activation, we cultured E14.5 SMGs from *Fgfr1^+/cKO^; Fgfr2^+/cKO^* controls, *Fgfr1^+/cKO^; Fgfr2^cKO/cKO^* and *Fgfr1^FCPG/cKO^; Fgfr2^cKO/cKO^* mutants in media supplemented with Mn^2+^ or in the presence of integrin-blocking antibodies. Changes in branching were analyzed at 18 hrs and 40 hrs of culture. At both time points, *Fgfr1^FCPG/cKO^; Fgfr2^cKO/cKO^* mutants showed similar extent of branching as *Fgfr1^+/cKO^; Fgfr2^cKO/cKO^* mutants, quantified by average curvature analysis and number of buds (Figure 7A, B; Supplementary Figure 7A). These results suggest that FGFR1 canonical signaling is not required for branching upon integrin signaling activation. We further evaluated changes in branching at 40 hrs upon Mn^2+^ treatment in *Fgfr2^FCPG/FCPG^* mutants. Compared to *Fgfr2^+/+^* and *Fgfr2^+/FCPG^* mutants, *Fgfr2^FCPG/FCPG^* mutants only showed a mild reduction in overall curvature (Figure 7C, Supplementary Figure 7B). These results suggest that both *Fgfr1^FCPG^* and *Fgfr2^FCPG^* signaling alleles can partially rescue branching defects during SMG development.

**Figure 7:**
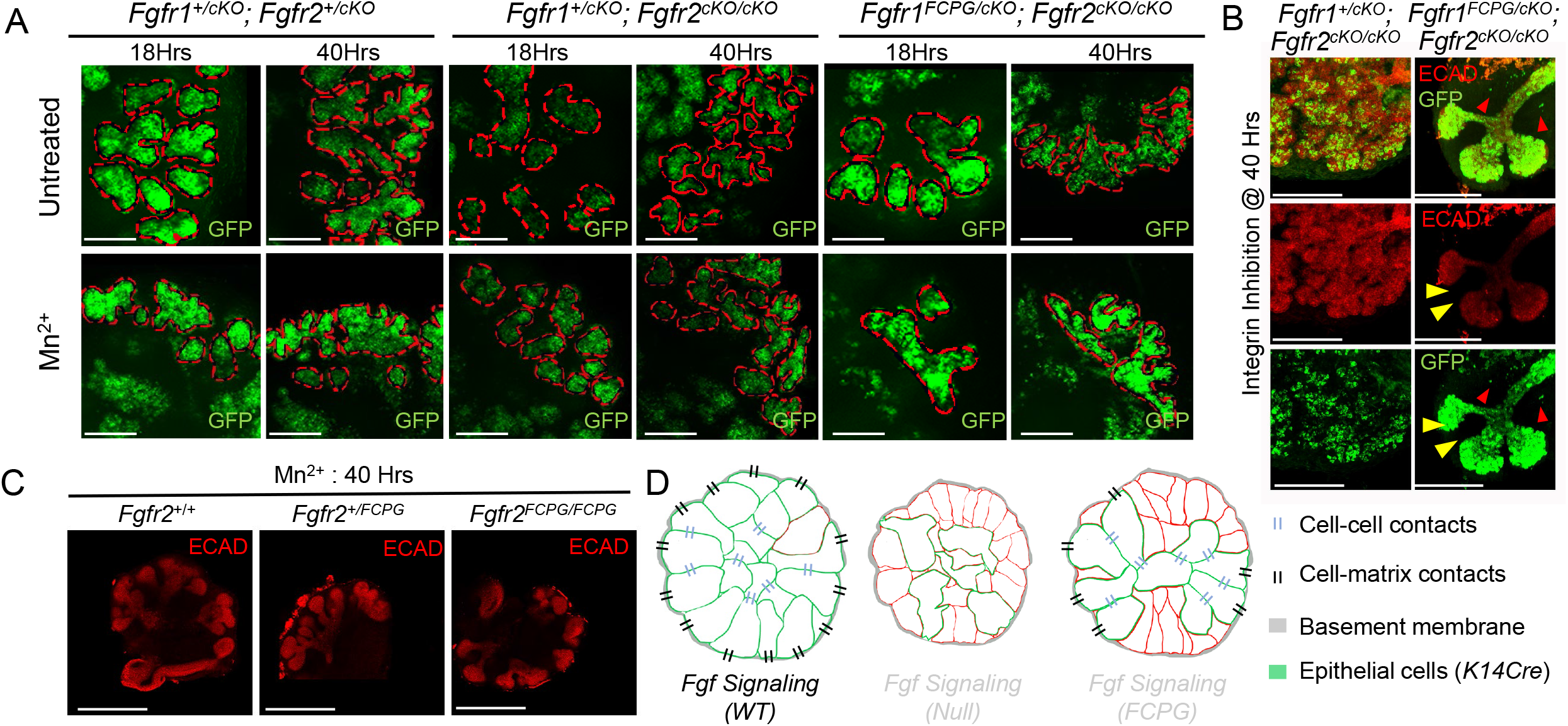
Integrin signaling activation rescues branching defects in *Fgfr1* and *Fgfr2* signaling mutants. (A) SMG explants (E14.5) from *Fgfr1* signaling mutants were cultured for 40 hrs in presence of Mn^2+^ to activate integrin signaling. Tissues were imaged at 18 hrs and 40 hrs. Representative images for respective genotypes are shown. Epithelial surfaces are depicted as dotted red lines. At 40 hrs, an increase in overall branching was observed across all mutants upon integrin signaling activation. Similar levels of branching were observed for all the mutants analyzed. Scale bars, 50 μm. (B) *Fgfr1^+/cKO^; Fgfr2^cKO/cKO^* (*n*=4) and *Fgfr1^FCPG/cKO^; Fgfr2^cKO/cKO^* mutant (*n=5*) SMG explants (E14.5) were cultured with a blocking antibody to inhibit integrinβ1 signaling. Tissues were analyzed 40 hrs post treatment. ECAD immunostaining was used to mark epithelial cells (yellow arrowhead). Only rarely were GFP^+^ cells observed outside epithelial population (red arrowhead). Scale bar 200 μm. (C) Integrin signaling activation in *Fgfr2* signaling mutants (at E14.5; Control *Fgfr2^+/FCPG^, Fgfr2^FCPG/FCPG^)* was carried out in culture for 40 hrs. After 40 hrs, ECAD immunostaining was used to mark the epithelial tissue. Representative images for respective genotype are shown. Median average curvature values were plotted as a bar graph for respective genotypes (Supplementary Figure 7). Overall, the extent of branching remained unchanged across all genotypes. Scale bars, 200 μm. (D) FGF signaling play a crucial role during SMG branching. Upon loss of FGF signaling, salivary gland epithelial cells fail to maintain strong cell-cell and cell-basement membrane interactions, resulting in defects in epithelial branching and reduction in the number of acini. Both branching and cell-cell contacts are partially restored by *Fgfr1^FCPG^* or *Fgfr2^FCPG^* mutants, indicating that non-canonical FGF signaling is important for maintaining strong cell-cell and cellbasement membrane interactions.

## DISCUSSION

Growth factors expressed in the salivary gland mesenchyme are known to signal to the epithelium to regulate epithelial morphogenesis, and previous genetic studies have shown that the FGF pathway is paramount in this process (Hoffman et al., 2002; Min et al., 1998; Ohuchi et al., 2000). Accordingly, we observed that *Fgfr1* and *Fgfr2* were highly expressed in the outer layer of the epithelial buds. To determine epithelial specific FGFR functions, we conditionally deleted *Fgfr1* and/or *Fgfr2* using a *K14Cre* driver. SMGs showed clear branching defects when either *Fgfr1* or *Fgfr2* activity was lost, with a more pronounced defect with loss of *Fgfr2.* Compound *Fgfr1/2 null* mutants developed the most severe defects, indicating that both receptors function coordinately during epithelial branching. However, the contribution of canonical intracellular signaling downstream of FGFs remained to be assessed. In this work, we tie FGF signaling to the ability of epithelial cells to adhere to the matrix and to each other, and show how FGF driven cellmatrix interactions regulate epithelial branching.

As RTKs, FGF receptors classically activate multiple signal transduction cascades upon receptor dimerization, including ERK1/2, PI3K, PLCγ, JNK, JAK/STAT and p38 (Brewer et al., 2016; Lanner and Rossant, 2010). ERK1/2 has widely been considered a hallmark downstream FGF signaling pathway (Brewer et al, 2016; Lanner and Rossant, 2010). However, ligand independent RAF-MEK-ERK activation, using a conditional constitutively active *Braf^CA^* allele, could not alleviate branching defects in compound *Fgfr1/2* null mutants. Conversely, *Fgfr1^FCPG^* and *Fgfr2^FCPG^* mutants, which broadly eliminate classic RTK signaling outputs (Ray et al., 2020), partially alleviated branching defects. In compound *Fgfr1/2 null* mutant background, restoration of a single *Fgfr1^FCPG^* allele could reconstitute terminal branching to a similar extent as a wild-type *Fgfr1* allele. Likewise, *Fgfr2^FCPG/FCPG^* mutants rescued by around half the SMG terminal branching defects, although it did not rescue SLG development. These results indicate that FGFs exert their functions in salivary glands, at least in part, through mechanisms beyond canonical RTK signaling.

Cell-cell and cell-matrix adhesion have been shown to play instructive roles in epithelial morphogenesis. In the salivary gland, the surface epithelial sheet establishes strong cell matrix adhesion to the BM, and inward folding of cells with weaker cell-cell adhesion promotes budding and accompanying clefting (Wang et al., 2021). We investigated cell adhesion in mutant salivary glands and organ culture, since FGFRs have been shown to interact with integrins and cadherins through their extracellular domain, which has not been altered in our most severe *Fgfr1^FCPG^* and *Fgfr2^FCPG^* signaling alleles (Endo et al., 2012; Geiger and Yamada, 2011; McQuade et al., 2006; Moser et al., 2009; Rapraeger et al., 1991; Yayon et al., 1991). Moreover, we previously observed that *Fgfr1^-/-^* and/or *Fgfr2^-/-^* primary neural crest cells failed to spread on fibronectin coated dishes in culture and to form extensive cell-cell contacts *in vivo,* but that both processes were restored in *Fgfr1* and/or *Fgfr2* signaling mutant backgrounds (Ray et al., 2020). Remarkably, we found similar behaviors with SMG epithelial cells, as *Fgfr1/2* null mutant acinar cells had compromised adhesion to the BM and failed to exhibit extensive cell-cell contacts. As in previous work, we observed that adhesion defects were rescued to a large extent by reintroducing *Fgfr1^FCPG^* and *Fgfr2^FCPG^* alleles to double mutant backgrounds, indicating that FGF regulation of cell adhesion operates by pathways independent of canonical RTK signaling (Figure 7D).

How FGF signaling works with cell adhesion pathways remains to be determined (Clark and Soriano, 2022; Ferguson et al., 2021). The fact that loss of FGFR affects cell-matrix adhesion, as shown here and in previous work (Ray et al., 2020), suggests that FGF signaling operates at some level upstream of cell-matrix adhesion pathways to regulate their activity. We observed that *Fgfr1^cKO/cKO^; Fgfr2^cKO/cKO^* mutant cells failed to populate the outer margins of the epithelial buds. Outer bud epithelial cells, which significantly co-express both *Fgfr1* and *Fgfr2*, normally show accumulation of integrinβ1. In addition, the integrity of the BM was affected by FGF signaling, as compound *Fgfr1/2 null* mutants develop ruptures of the BM *in vivo*, with some cells extruding into the surrounding mesenchyme. Inhibiting integrin activation in culture led to a similar phenotype. It has been suggested that interactions with integrins might seed laminin polymerization to initiate BM assembly (Yurchenco and Patton, 2009). Therefore, reduced integrin activation might lead to a reduction in BM formation and thus be responsible for the BM ruptures. Conversely, activating integrin signaling in an *ex vivo* culture model resulted in a significant increase in SMG branching, but was unable to rescue the extensive defects in conditional *Fgfr1/2* null mutant cultures, which still exhibited defects in the BM. Under similar conditions, reintroducing a single *Fgfr1^FCPG^* mutant allele, and to a lesser extent a similar *Fgfr2^FCPG^* copy, could rescue *Fgfr1/2* null mutant defects.

Alternatively, it is possible that *Fgfr1^cKO/cKO^; Fgfr2^cKO/cKO^* mutant cells can punch through the basement membrane, similar to metastatic cancer cells or *C.elegans* anchor cells, due to weakened integrin signaling but also possibly improper cadherin switching where FGF has been shown to play a critical role (Sun et al. 1999; Ciruna and Rossant 2001; Nieto et al. 2016). The escape of these cells might also explain why we find fewer *K14Cre* lineage GFP^+^ cells in mutant salivary glands, since we failed to observe a defect in cell proliferation or cell death at E14.5 despite the known roles of FGF signaling in mediating these processes. The effect that we have observed upon loss of FGF signaling on ECAD expression ties into previous investigations, as FGF signaling has been shown to affect E-cadherin localization in multiple contexts, including the mural trophectoderm (Kurowski et al. 2019), *Drosophila* mesoderm (Sun and Stathopoulos 2018), and zebrafish cardiomyocytes (Rasouli et al. 2018). Taken together, our observations suggest that non-canonical FGF signaling plays a significant role in regulating cell-cell and cell-matrix interactions. Further investigations will shed light into how these interactions come into play to remodel the BM and promote branching of the epithelial tissue during salivary gland branching morphogenesis.

## MATERIALS AND METHODS

### Mouse strains

All animal experiments were approved by the Institutional Animal Care and Use Committee at the Icahn School of Medicine at Mount Sinai. *Fgfr1^Lox^ (Fgfr1^tm5.1Sor^), Fgfr2^Lox^ (Fgfr2^tm1.1Sor^), Fgfr1^FCPG^ (Fgfr1^tm10.1Sor^), Fgfr2^FCPG^ (Fgfr2^tm8.1Sor^), Fgfr1-GFP (Fgfr1^tm12.1Sor^)* and *Fgfr2-mCherry* (*Fgfr2^tm2.1Sor^*) were previously described (Hoch and Soriano, 2006; Molotkov et al., 2017). (*Cg)-Braf^tm1Mmcm^*/J, a conditional constitutive active *Braf* allele is referred in the text as *Braf^CA^*, has been described previously (Dankort et al., 2007). *Tg(KRT14-cre)Smr, Gt(ROSA)26Sor^tm1Sor^* and *Gt(ROSA)26Sor^tm4(ACTB-tdTomato,-EGFP)Luo^* are referred to in the text as *K14-Cre, ROSA26^LacZ^* and *ROSA26^mT/mG^*, respectively (Andl et al., 2004; Muzumdar et al., 2007; Soriano, 1999). All lines were maintained on a 129S4 co-isogenic background, except for *Braf^CA^* that was used in crosses after four generations of backcross to 129S4. Successful mating for all timed pregnancies were assessed by the presence of a vaginal plug and designated embryonic day 0.5 (E0.5).

### Whole mount immunostaining

The extent of branching in the developing salivary glands was analyzed on whole mounts. Intact salivary glands (stromal mesenchyme + epithelium) were harvested for wild-type or different *Fgfr1/2* conditional/signaling mutants in 1 x PBS. Tissues were washed and fixed overnight in 4% paraformaldehyde in PBS (4%PFA/PBS) at 4°C. Fixed tissues were washed three times in PBS followed by permeabilization in 1 x PBT (PBS supplemented with 0.5% TritonX100) for the following 2 days with intermittent changes of fresh PBT. 5%BSA in PBT was used for blocking prior to antibody incubations for another 2 days at 4°C. This was followed by incubation with appropriate dilution of primary antibodies/blocking solution for whole-mount immunostaining for 5 days. Salivary glands were washed in PBT multiple times for the next 2 days before further incubations with appropriate secondary antibody in PBT, followed by counterstaining with DAPI. After washing, tissues were gradually dehydrated to 100% methanol in a gradient of increasing methanol concentration followed by clearing by multiple changes of (1:3) benzyl alcohol: benzyl benzoate (BABB). Whole mount imaging was carried out on a Leica TCS SP8 confocal microscope. Imaris 9.8.0 (Oxford Instruments) was used to import Z-stack image sequences and compiled to create 3D images. The number of terminal buds was counted for wild-type or different *Fgfr1/2* conditional mutants from 3D rendered images across XY rotational axes using Imaris 9.8.0.

### Tissue processing for sections

Salivary glands were harvested at E14.5, E15.5 and E18.5 for various *Fgfr1/2* conditional mutants in 1 x PBS. Tissues were fixed in 4% PFA/PBS overnight followed by washes in 1 x PBS. Fixed tissues were next incubated in increasing sucrose concentrations gradients (10–30%) in PBS and finally embedded in OCT. 10 μm frozen sections were cut using a cryostat (Leica) and stored at −20°C for analysis. Prior to immunofluorescence, slides were washed 3 times for 5 mins each in 1 x PBT, prior to blocking in 5% BSA/PBT for 1 hour. Tissues were next incubated with primary antibody at appropriate dilutions.

For LacZ staining, tissues were fixed in 2% formaldehyde and 0.2% glutaraldehyde in PBS for 10 mins followed by OCT embedding. 10 μm sections were generated. During detection, tissues were rehydrated by washing in 1 x PBS. Tissues were next washed in 5 mM potassium ferricyanide, 5 mM potassium ferrocyanide, 2 mM MgCI2, 0.01% sodium deoxycholate, and 0.02% Nonidet P-40 (NP-40) in PBS, before adding 1 mg/ml X-Gal in the same solution to detect lacZ activity.

For immunofluorescence, primary antibody incubations were carried out overnight at 4°C. Next, slides were washed 3 times 5 mins each in PBT before incubating with appropriate secondary antibodies for 5 hrs at room temperature. The primary antibodies and dilutions used were: GFP (1:500; AvesLab, #GFP-1020), mCHERRY (1:100, Abcam, #ab167453), ECAD (1:200, CST, #3195S), LAMININ (1:200, MilliporeSigma, #L9393), Integrinβ1 (1:100; Abcam, #ab30394), Activated Integrinβ1 (1:100, MilliporeSigma, #MAB2079-AF647, direct immunofluorescence). All secondary antibodies were used at 1:5,000 dilutions as follows: Alexa Fluor^®^ 488 AffiniPure Donkey Anti-Chicken IgY (IgG) (H+L) (Jackson ImmunoResearch Laboratories; #703-545-155); Donkey anti-Rabbit IgG (H+L), Alexa Fluor 488; Alexa Fluor 546; Alexa Fluor 647; Donkey anti-Mouse IgG (H+L), Alexa Fluor 488, Alexa Fluor 546; Alexa Fluor 647 (Thermo Fisher Scientific #A-21206; #A-10040; #A-31573; #A-21202; #A-10036; #A-31571). DAPI (1 μg/ml: Life Technologies, D1306) was used for nuclei staining together with secondary antibodies.

### Cell proliferation assay

14.5 dpc pregnant females were injected intraperitoneally with 100 mg/kg body weight of EdU. Salivary glands were harvested from embryos 1hour after EdU injection. Fixed tissues were processed and 10 μm sections were generated. EdU detection was carried out as per manufacturer’s instruction using the Click-iT EdU Cell Proliferation Kit (Thermo Fisher; #C10340) on sections.

### TUNEL assay

Sections generated from E14.5 salivary glands were rehydrated in PBS, followed by post-fixation in 4% PFA for 10 mins. Tissues were next washed in 1 x PBS to remove traces of PFA followed by washes in PBT (PBS + 0.1%Triton X100). Next, tissues were blocked in 5%BSA/PBT. The TUNEL In Situ Cell Death Detection Kit TMR red (MilliporeSigma; #12156792910) was used to detect cell death.

### Curvature analysis

Salivary gland epithelial tissues were labeled in whole mounts followed by tissue clearing and confocal microscopy imaging. Optical sections were generated. Curvature features were extracted from tissue sections immunostained for Laminin that mark BM. The curvature of the peripheral epithelial cell layer at the epithelial boundaries or from the BM marked by Laminin, were analyzed using the Kappa plugin in the FIJI implementation of ImageJ (Mary and Brouhard, 2019). Similarly, curvature features were extracted from explant cultures, either from the images collected at different timepoints or from ECAD immunostained fixed tissues. Closed B-Spline curves were traced, and the average curvature was estimated for SMGs. The data (median with 95%CI) was plotted as a bar graph or violin plots. Cell-BM interaction maps were generated using Imaris 9.8.0 (Oxford Instrument). 3D maps were generated using COLOC function to give a visual estimate of cell-BM interactions.

### Isolation and culture of Salivary gland

Mouse salivary glands were isolated at E14.5 by decapitating embryos with fine scissors. In a fresh plate containing PBS, a scalpel was used to slice in between the lower and the upper jaw to separate the mandible and tongue from the rest of the head. The salivary glands, sandwiched between the base of the tongue and the lower jaw, were next transferred to a fresh plate containing PBS on a dissection scope. The tissue was positioned with the tongue facing the top. A pair of forceps was used to gently lift the tongue and subsequently two salivary glands on either side were removed and transferred to a fresh plate with DMEM/F-12 (Thermo Fisher, 11039047) medium prior to culture.

Isolated salivary glands were cultured in Ibidi 8-well μ-slides. Upon coating with Corning GFR Matrigel (1:5 dilution; Thermo Fisher #CB-40230), salivary glands were laid out avoiding drying and overlaid with organ culture medium containing DMEM/F-12 supplemented with 150 μg/mL vitamin C (MilliporeSigma, A7506), 2XITS and 1 × PenStrep (100 units/mL penicillin, 100 μg/mL streptomycin; Thermo Fisher, 15140163). Tissues were incubated at 37°C in 5% CO_2_ for 40 hours.

Organ culture medium was supplemented with 50 μM MnCl2 to enhance integrin-mediated cellmatrix adhesion signaling functions. To block β1-integrin functions, we either used rat monoclonal (mAb13) β1-integrin blocking antibodies at 100 μg/mL (MilliporeSigma; # MABT821) or mouse monoclonal (P5D2) β1-integrin blocking antibodies at 50 μg/mL (Abcam; ab24693).

### Statistical analysis

Unpaired two-tailed Student’s *t*-test were used to obtain *P* values. The error bars represent standard deviations of means for all the graphs except for the average curvature analysis. P-values are summarized as “ns” (P>0.05), “*” (P<0.05), “**” (P<0.01), “***” (P<0.001) and “****” (P<0.0001).

For all graphs where average curvatures were analyzed, median average curvature were either represented as violin plots or as bar graphs. The error bars represent 95% confidence intervals.

## Acknowledgments

We thank our laboratory colleagues, Rob Krauss, Ali May and Checco Ramirez for helpful discussions and critical comments on the manuscript. We thank Yeifei Sun for help on analyzing scRNA seq data, and the Microscopy and Advanced Imaging Core for assistance and suggestions on data analysis. This work was supported by grant DE022778 from the National Institute of Dental and Craniofacial Research to P.S.

---

**Supplementary Figure 1.**
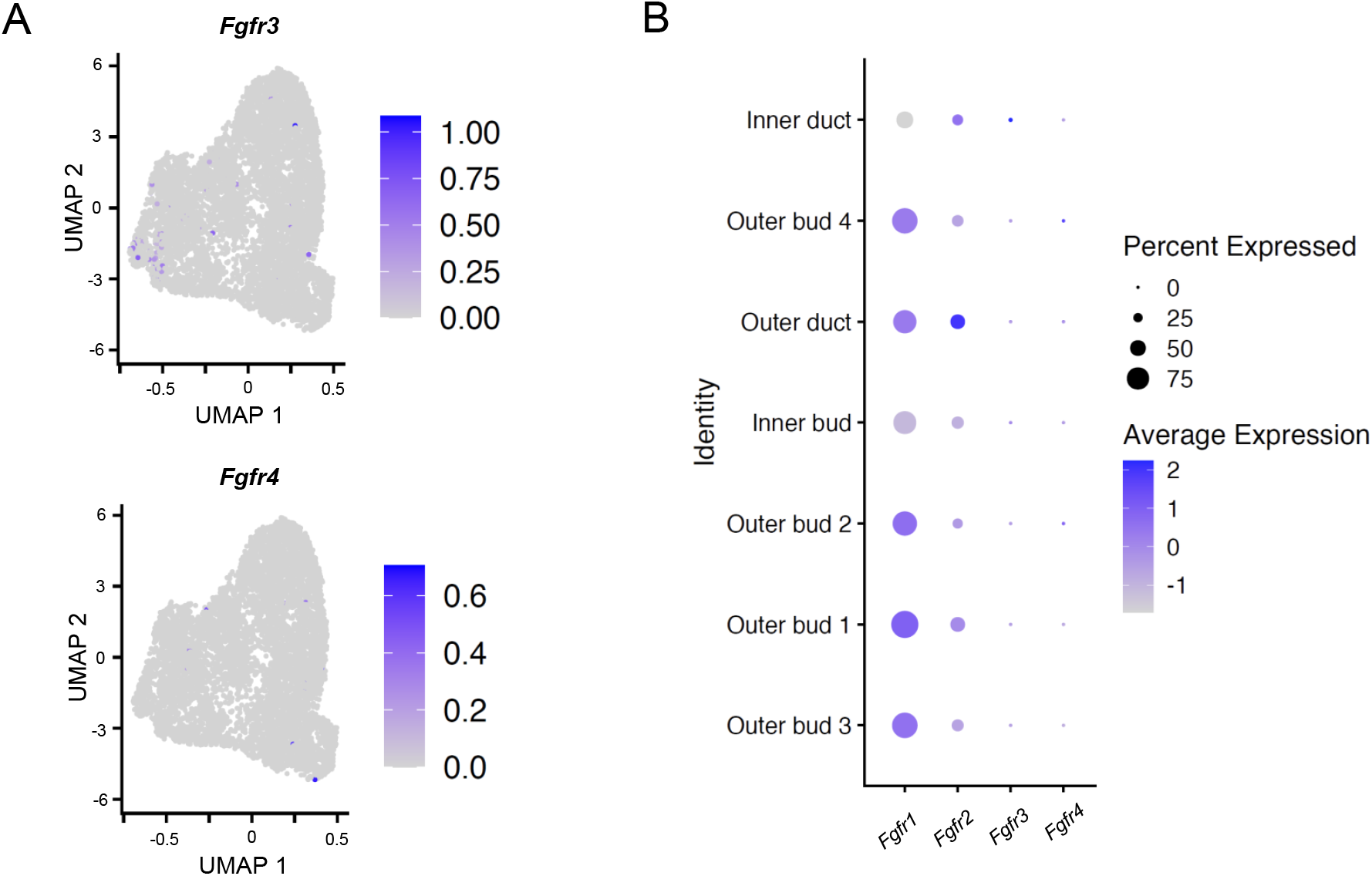
(related to Figure 1)

**Supplementary Figure 2.**
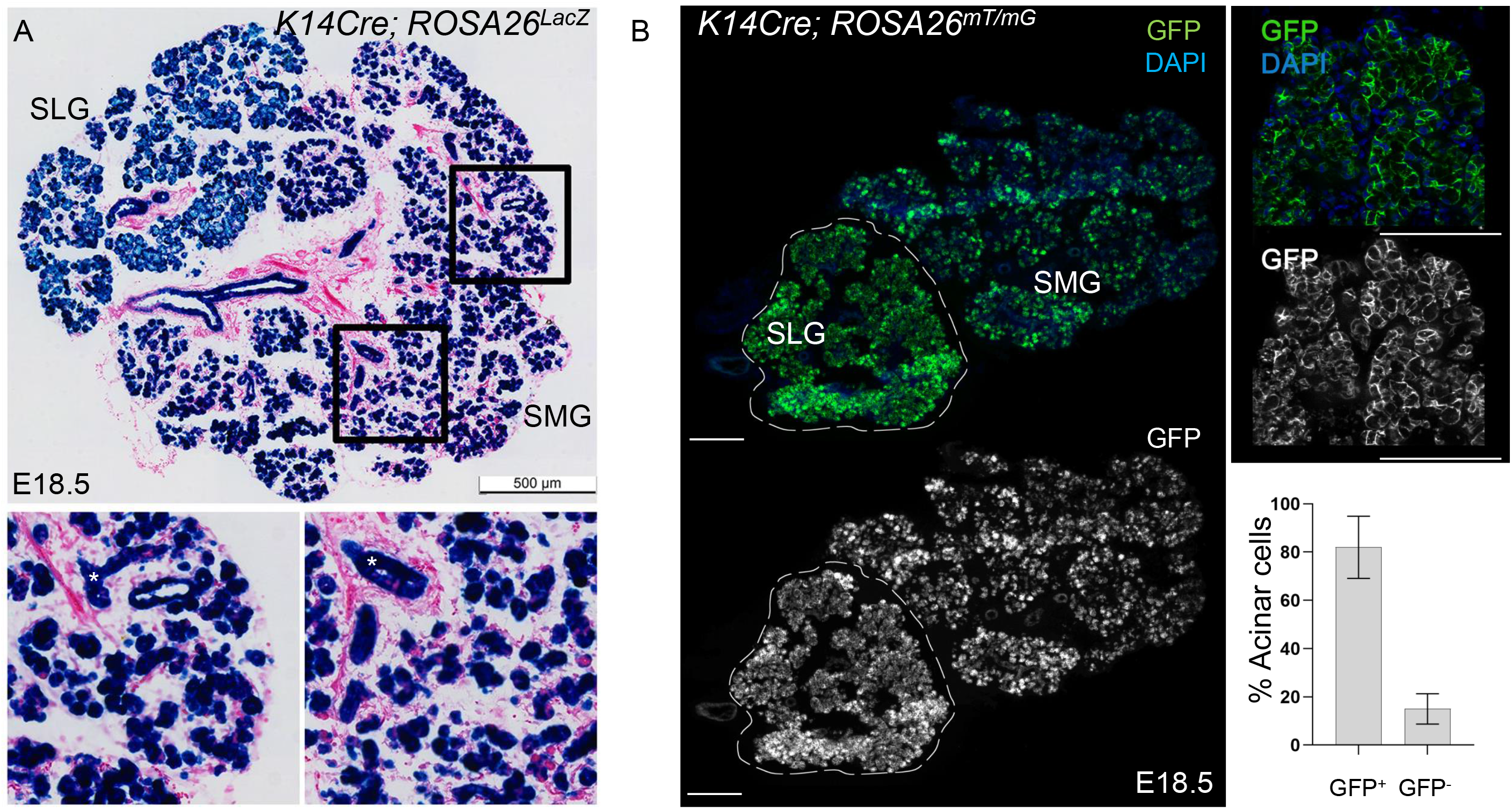
A,B (related to Figure 2)

**Supplementary Figure 3.**
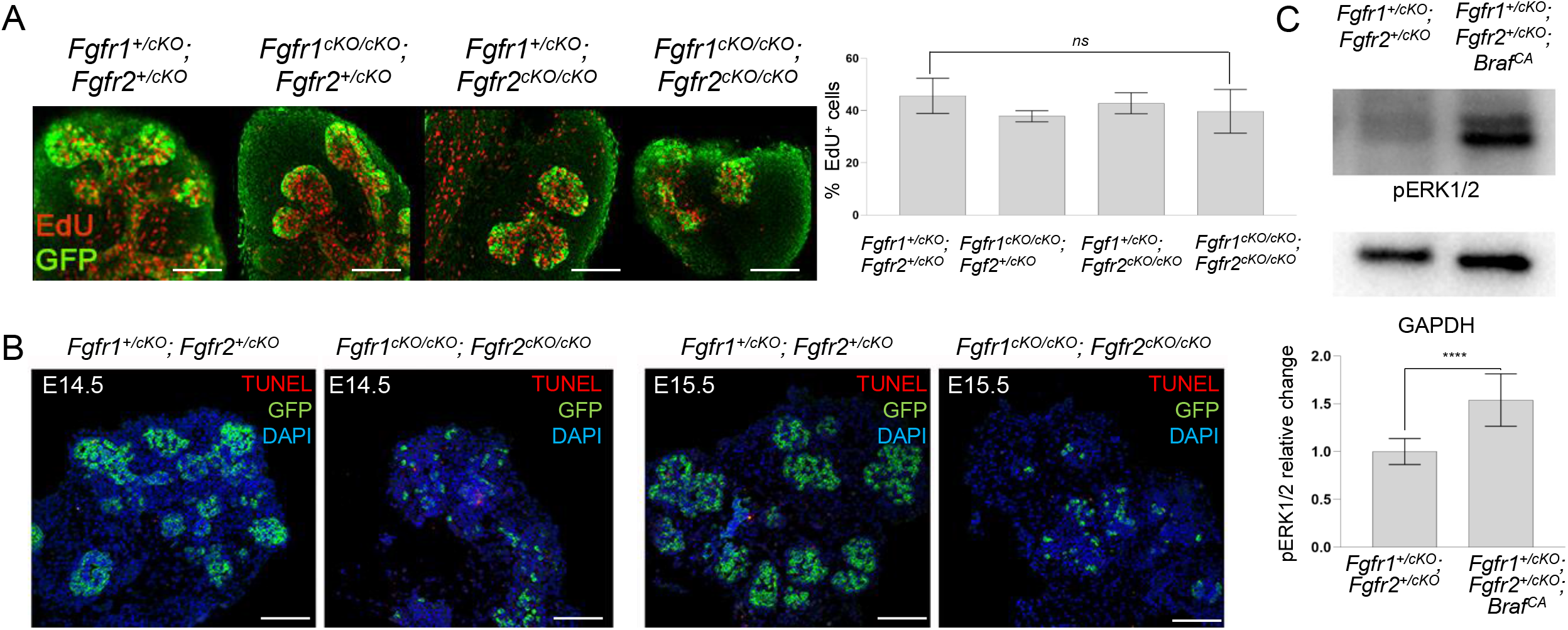
(related to Figure 3)

**Supplementary Figure 4.**
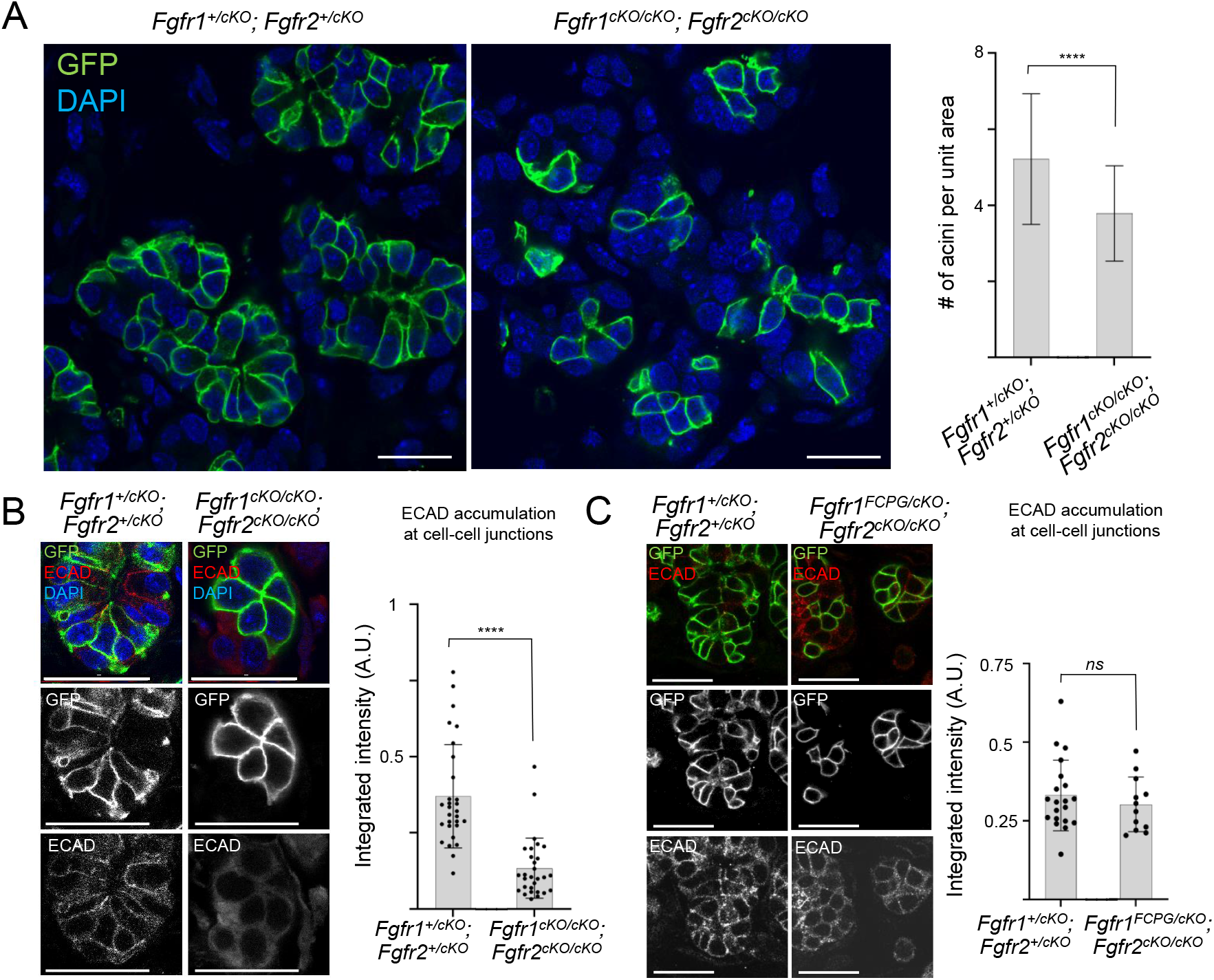
(related to Figure 4)

**Supplementary Figure 5.**
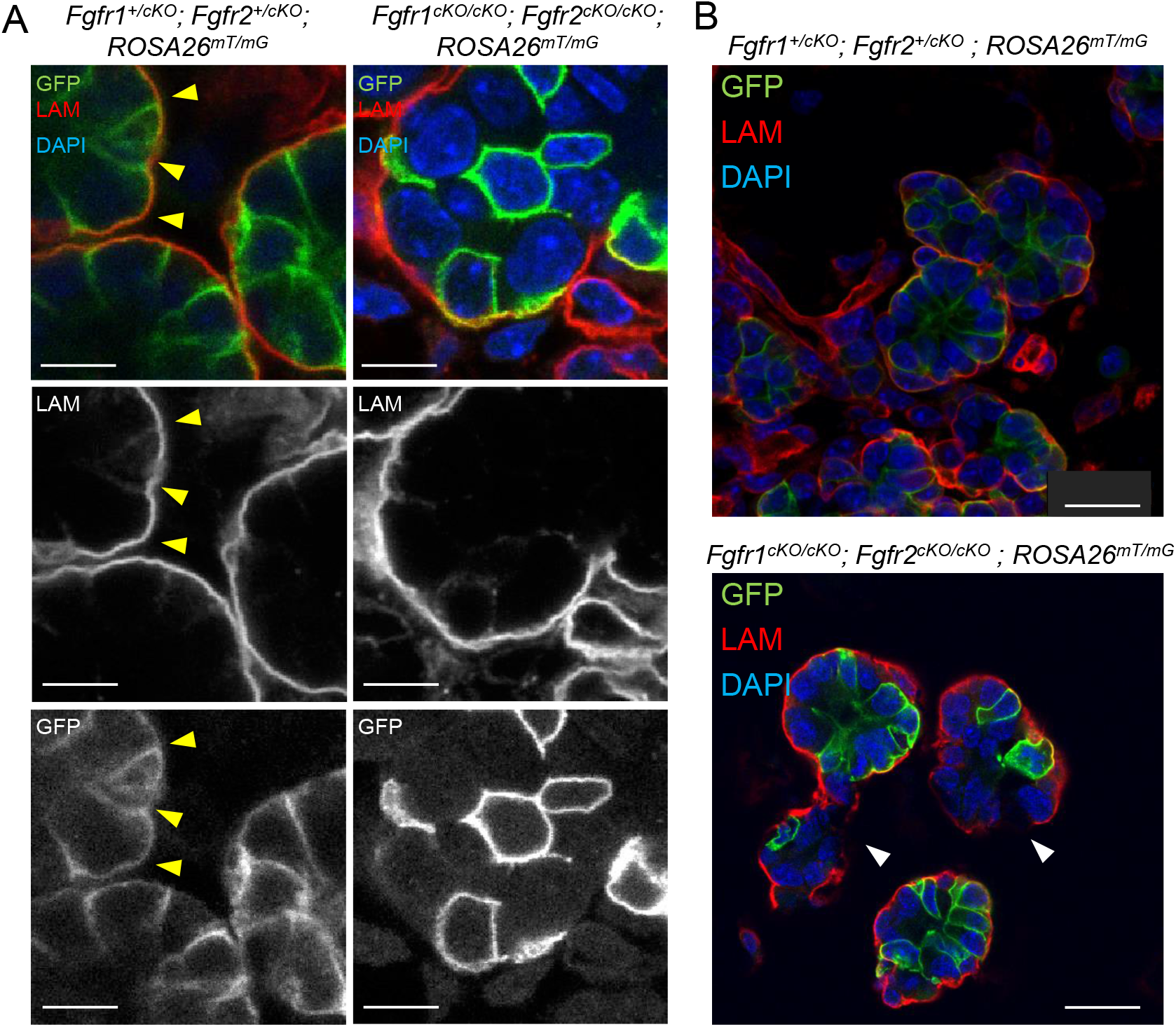
(related to Figure 5)

**Supplementary Figure 6.**
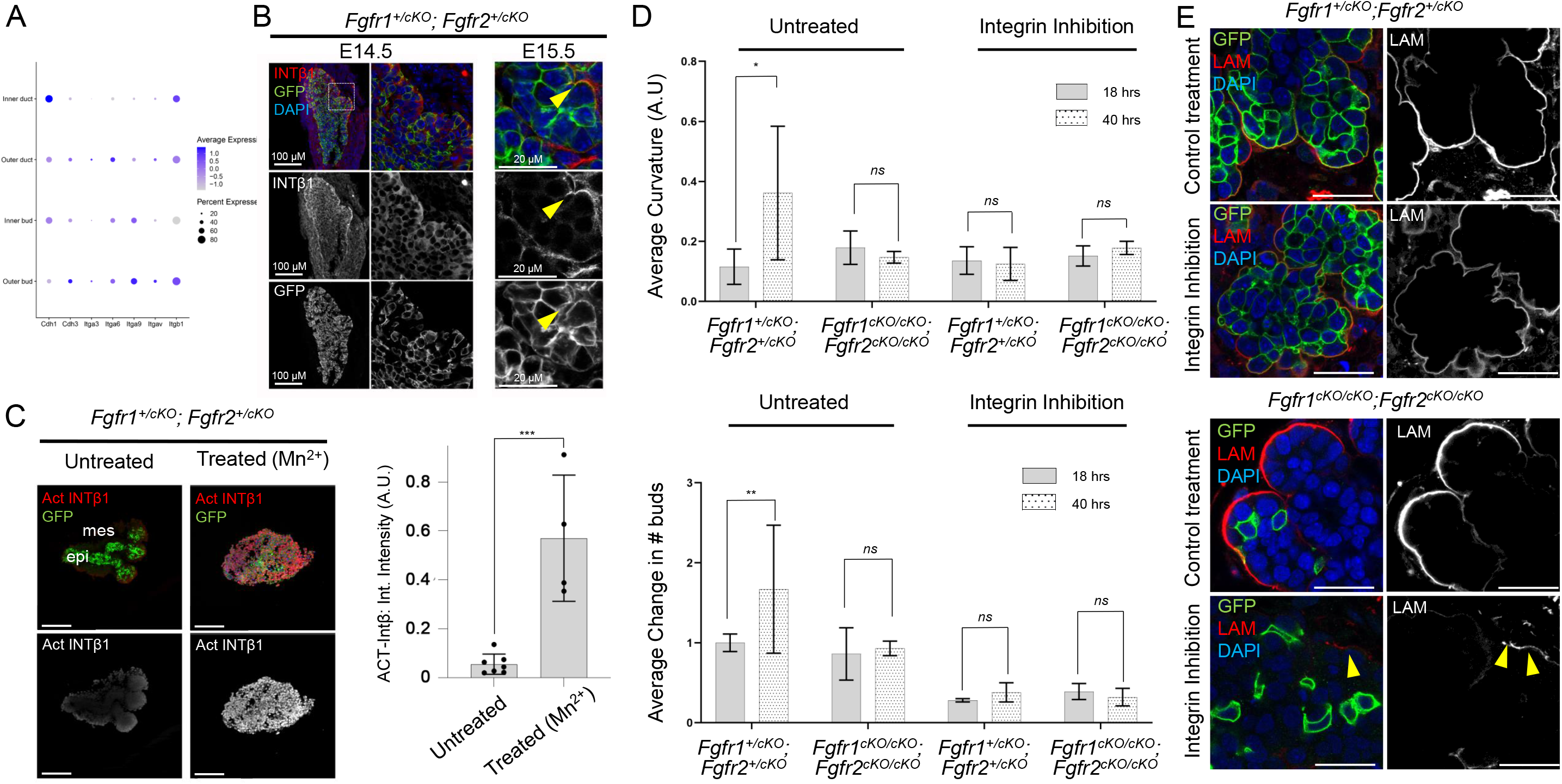
(related to Figure 6)

**Supplementary Figure 7.**
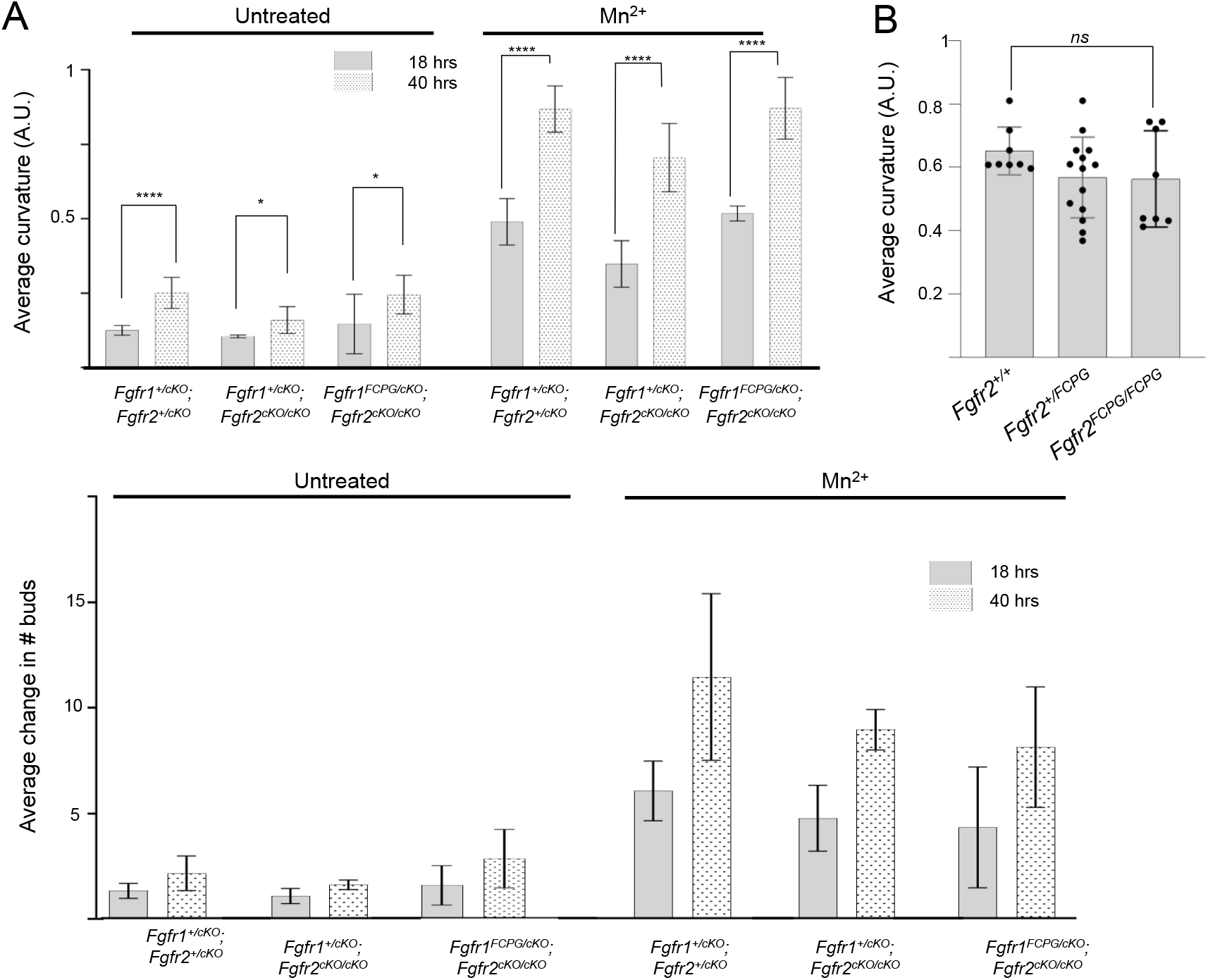
(related to Figure 7)

## References

Andl, T., Ahn, K., Kairo, A., Chu, E.Y., Wine-Lee, L., Reddy, S.T., Croft, N.J., Cebra-Thomas, J.A., Metzger, D., Chambon, P., et al. (2004). Epithelial Bmpr1a regulates differentiation and proliferation in postnatal hair follicles and is essential for tooth development. Development 131, 2257–2268.

Bazzoni, G., Shih, D.T., Buck, C.A., and Hemler, M.E. (1995). Monoclonal antibody 9EG7 defines a novel beta 1 integrin epitope induced by soluble ligand and manganese, but inhibited by calcium. J Biol Chem 270, 25570–25577.

Brewer, J.R., Mazot, P., and Soriano, P. (2016). Genetic insights into the mechanisms of Fgf signaling. Genes Dev 30, 751–771.

Brewer, J.R., Molotkov, A., Mazot, P., Hoch, R.V., and Soriano, P. (2015). Fgfr1 regulates development through the combinatorial use of signaling proteins. Genes Dev 29, 1863–1874.

Celli, G., LaRochelle, W.J., Mackem, S., Sharp, R., and Merlino, G. (1998). Soluble dominant-negative receptor uncovers essential roles for fibroblast growth factors in multi-organ induction and patterning. EMBO J 17, 1642–1655.

Chatzeli, L., Gaete, M., and Tucker, A.S. (2017). Fgf10 and Sox9 are essential for the establishment of distal progenitor cells during mouse salivary gland development. Development 144, 2294–2305.

Clark, J.F., and Soriano, P.M. (2022). Pulling back the curtain: The hidden functions of receptor tyrosine kinases in development. Curr Top Dev Biol 149, 123–152.

Costantini, F., and Kopan, R. (2010). Patterning a complex organ: branching morphogenesis and nephron segmentation in kidney development. Dev Cell 18, 698–712.

Dankort, D., Filenova, E., Collado, M., Serrano, M., Jones, K., and McMahon, M. (2007). A new mouse model to explore the initiation, progression, and therapy of BRAFV600E-induced lung tumors. Genes Dev 21, 379–384.

De Moerlooze, L., Spencer-Dene, B., Revest, J.M., Hajihosseini, M., Rosewell, I., and Dickson, C. (2000). An important role for the IIIb isoform of fibroblast growth factor receptor 2 (FGFR2) in mesenchymal-epithelial signalling during mouse organogenesis. Development 127, 483–492.

Doherty, P., and Walsh, F.S. (1996). CAM-FGF Receptor Interactions: A Model for Axonal Growth. Mol Cell Neurosci 8, 99–111.

Endo, Y., Ishiwata-Endo, H., and Yamada, K.M. (2012). Extracellular matrix protein anosmin promotes neural crest formation and regulates FGF, BMP, and WNT activities. Dev Cell 23, 305–316.

Entesarian, M., Dahlqvist, J., Shashi, V., Stanley, C.S., Falahat, B., Reardon, W., and Dahl, N. (2007). FGF10 missense mutations in aplasia of lacrimal and salivary glands (ALSG). Eur J Hum Genet 15, 379–382.

Entesarian, M., Matsson, H., Klar, J., Bergendal, B., Olson, L., Arakaki, R., Hayashi, Y., Ohuchi, H., Falahat, B., Bolstad, A.I., et al. (2005). Mutations in the gene encoding fibroblast growth factor 10 are associated with aplasia of lacrimal and salivary glands. Nat Genet 37, 125–127.

Ewald, A.J., Brenot, A., Duong, M., Chan, B.S., and Werb, Z. (2008). Collective epithelial migration and cell rearrangements drive mammary branching morphogenesis. Dev Cell 14, 570–581.

Ferguson, H.R., Smith, M.P., and Francavilla, C. (2021). Fibroblast Growth Factor Receptors (FGFRs) and Noncanonical Partners in Cancer Signaling. Cells 10.

Francavilla, C., Loeffler, S., Piccini, D., Kren, A., Christofori, G., and Cavallaro, U. (2007). Neural cell adhesion molecule regulates the cellular response to fibroblast growth factor. J Cell Sci 120, 4388–4394.

Geiger, B., and Yamada, K.M. (2011). Molecular architecture and function of matrix adhesions. Cold Spring Harb Perspect Biol 3.

Guo, L., Degenstein, L., and Fuchs, E. (1996). Keratinocyte growth factor is required for hair development but not for wound healing. Genes Dev 10, 165–175.

Harunaga, J.S., Doyle, A.D., and Yamada, K.M. (2014). Local and global dynamics of the basement membrane during branching morphogenesis require protease activity and actomyosin contractility. Dev Biol 394, 197–205.

Hoch, R.V., and Soriano, P. (2006). Context-specific requirements for Fgfr1 signaling through Frs2 and Frs3 during mouse development. Development 133, 663–673.

Hoffman, M.P., Kidder, B.L., Steinberg, Z.L., Lakhani, S., Ho, S., Kleinman, H.K., and Larsen, M. (2002). Gene expression profiles of mouse submandibular gland development: FGFR1 regulates branching morphogenesis in vitro through BMP-and FGF-dependent mechanisms. Development 129, 5767–5778.

Jaskoll, T., Abichaker, G., Witcher, D., Sala, F.G., Bellusci, S., Hajihosseini, M.K., and Melnick, M. (2005). FGF10/FGFR2b signaling plays essential roles during in vivo embryonic submandibular salivary gland morphogenesis. BMC Dev Biol 5, 11.

Jaskoll, T., Witcher, D., Toreno, L., Bringas, P., Moon, A.M., and Melnick, M. (2004). FGF8 dose-dependent regulation of embryonic submandibular salivary gland morphogenesis. Dev Biol 268, 457–469.

Kadoya, Y., Kadoya, K., Durbeej, M., Holmvall, K., Sorokin, L., and Ekblom, P. (1995). Antibodies against domain E3 of laminin-1 and integrin alpha 6 subunit perturb branching epithelial morphogenesis of submandibular gland, but by different modes. J Cell Biol 129, 521–534.

Kadoya, Y., and Yamashina, S. (1993). Distribution of alpha 6 integrin subunit in developing mouse submandibular gland. J Histochem Cytochem 41, 1707–1714.

Kashimata, M., and Gresik, E.W. (1997). Epidermal growth factor system is a physiological regulator of development of the mouse fetal submandibular gland and regulates expression of the alpha6-integrin subunit. Dev Dyn 208, 149–161.

Kashimata, M., Sayeed, S., Ka, A., Onetti-Muda, A., Sakagami, H., Faraggiana, T., and Gresik, E.W. (2000). The ERK-1/2 signaling pathway is involved in the stimulation of branching morphogenesis of fetal mouse submandibular glands by EGF. Dev Biol 220, 183–196.

Lanner, F., and Rossant, J. (2010). The role of FGF/Erk signaling in pluripotent cells. Development 137, 3351–3360.

Larsen, M., Hoffman, M.P., Sakai, T., Neibaur, J.C., Mitchell, J.M., and Yamada, K.M. (2003). Role of PI 3-kinase and PIP3 in submandibular gland branching morphogenesis. Dev Biol 255, 178–191.

Larsen, M., Yamada, K.M., and Musselmann, K. (2010). Systems analysis of salivary gland development and disease. Wiley Interdiscip Rev Syst Biol Med 2, 670–682.

Latko, M., Czyrek, A., Porebska, N., Kucinska, M., Otlewski, J., Zakrzewska, M., and Opalinski, L. (2019). Cross-Talk between Fibroblast Growth Factor Receptors and Other Cell Surface Proteins. Cells 8.

Mailleux, A.A., Spencer-Dene, B., Dillon, C., Ndiaye, D., Savona-Baron, C., Itoh, N., Kato, S., Dickson, C., Thiery, J.P., and Bellusci, S. (2002). Role of FGF10/FGFR2b signaling during mammary gland development in the mouse embryo. Development 129, 53–60.

Makarenkova, H.P., Hoffman, M.P., Beenken, A., Eliseenkova, A.V., Meech, R., Tsau, C., Patel, V.N., Lang, R.A., and Mohammadi, M. (2009). Differential interactions of FGFs with heparan sulfate control gradient formation and branching morphogenesis. Sci Signal 2, ra55.

Makarenkova, H.P., Ito, M., Govindarajan, V., Faber, S.C., Sun, L., McMahon, G., Overbeek, P.A., and Lang, R.A. (2000). FGF10 is an inducer and Pax6 a competence factor for lacrimal gland development. Development 127, 2563–2572.

Mary, H., and Brouhard, G.J. (2019). Kappa (K): Analysis of Curvature in Biological Image Data using B-splines. bioRxiv, 852772.

May, A.J., Headon, D., Rice, D.P., Noble, A., and Tucker, A.S. (2016). FGF and EDA pathways control initiation and branching of distinct subsets of developing nasal glands. Dev Biol 419, 348–356.

May, A.J., Teshima, T.H.N., Noble, A., and Tucker, A.S. (2019). FGF10 is an essential regulator of tracheal submucosal gland morphogenesis. Dev Biol 451, 158–166.

McQuade, K.J., Beauvais, D.M., Burbach, B.J., and Rapraeger, A.C. (2006). Syndecan-1 regulates alphavbeta5 integrin activity in B82L fibroblasts. J Cell Sci 119, 2445–2456.

Milunsky, J.M., Zhao, G., Maher, T.A., Colby, R., and Everman, D.B. (2006). LADD syndrome is caused by FGF10 mutations. Clin Genet 69, 349–354.

Min, H., Danilenko, D.M., Scully, S.A., Bolon, B., Ring, B.D., Tarpley, J.E., DeRose, M., and Simonet, W.S. (1998). Fgf-10 is required for both limb and lung development and exhibits striking functional similarity to Drosophila branchless. Genes Dev 12, 3156–3161.

Miner, J.H., and Yurchenco, P.D. (2004). Laminin functions in tissue morphogenesis. Annu Rev Cell Dev Biol 20, 255–284.

Molotkov, A., Mazot, P., Brewer, J.R., Cinalli, R.M., and Soriano, P. (2017). Distinct Requirements for FGFR1 and FGFR2 in Primitive Endoderm Development and Exit from Pluripotency. Dev Cell 41, 511–526 e514.

Moser, M., Legate, K.R., Zent, R., and Fassler, R. (2009). The tail of integrins, talin, and kindlins. Science 324, 895–899.

Moskwa, N., Mahmood, A., Nelson, D.A., Altrieth, A.L., Forni, P.E., and Larsen, M. (2022). Singlecell RNA sequencing reveals PDFGRalpha+ stromal cell subpopulations that promote proacinar cell differentiation in embryonic salivary gland organoids. Development 149.

Muzumdar, M.D., Tasic, B., Miyamichi, K., Li, L., and Luo, L. (2007). A global double-fluorescent Cre reporter mouse. Genesis 45, 593–605.

Nguyen, T., and Mege, R.M. (2016). N-Cadherin and Fibroblast Growth Factor Receptors crosstalk in the control of developmental and cancer cell migrations. Eur J Cell Biol 95, 415–426.

Nie, X., Luukko, K., and Kettunen, P. (2006). FGF signalling in craniofacial development and developmental disorders. Oral Dis 12, 102–111.

Nogawa, H., and Takahashi, Y. (1991). Substitution for mesenchyme by basement-membrane-like substratum and epidermal growth factor in inducing branching morphogenesis of mouse salivary epithelium. Development 112, 855–861.

Ohuchi, H., Hori, Y., Yamasaki, M., Harada, H., Sekine, K., Kato, S., and Itoh, N. (2000). FGF10 acts as a major ligand for FGF receptor 2 IIIb in mouse multi-organ development. Biochem Biophys Res Commun 277, 643–649.

Ornitz, D.M., and Itoh, N. (2022). New developments in the biology of fibroblast growth factors. WIREs Mech Dis, e1549.

Patel, V.N., Rebustini, I.T., and Hoffman, M.P. (2006). Salivary gland branching morphogenesis. Differentiation 74, 349–364.

Rapraeger, A.C., Krufka, A., and Olwin, B.B. (1991). Requirement of heparan sulfate for bFGF-mediated fibroblast growth and myoblast differentiation. Science 252, 1705–1708.

Ray, A.T., Mazot, P., Brewer, J.R., Catela, C., Dinsmore, C.J., and Soriano, P. (2020). FGF signaling regulates development by processes beyond canonical pathways. Genes Dev 34, 1735–1752.

Rebustini, I.T., Patel, V.N., Stewart, J.S., Layvey, A., Georges-Labouesse, E., Miner, J.H., and Hoffman, M.P. (2007). Laminin alpha5 is necessary for submandibular gland epithelial morphogenesis and influences FGFR expression through beta1 integrin signaling. Dev Biol 308, 15–29.

Sakakura, T., Nishizuka, Y., and Dawe, C.J. (1976). Mesenchyme-dependent morphogenesis and epithelium-specific cytodifferentiation in mouse mammary gland. Science 194, 1439–1441.

Sanchez-Heras, E., Howell, F.V., Williams, G., and Doherty, P. (2006). The fibroblast growth factor receptor acid box is essential for interactions with N-cadherin and all of the major isoforms of neural cell adhesion molecule. J Biol Chem 281, 35208–35216.

Shams, I., Rohmann, E., Eswarakumar, V.P., Lew, E.D., Yuzawa, S., Wollnik, B., Schlessinger, J., and Lax, I. (2007). Lacrimo-auriculo-dento-digital syndrome is caused by reduced activity of the fibroblast growth factor 10 (FGF10)-FGF receptor 2 signaling pathway. Mol Cell Biol 27, 6903–6912.

Soriano, P. (1999). Generalized lacZ expression with the ROSA26 Cre reporter strain. Nat Genet 21, 70–71.

Steinberg, Z., Myers, C., Heim, V.M., Lathrop, C.A., Rebustini, I.T., Stewart, J.S., Larsen, M., and Hoffman, M.P. (2005). FGFR2b signaling regulates ex vivo submandibular gland epithelial cell proliferation and branching morphogenesis. Development 132, 1223–1234.

Takahashi, Y., and Nogawa, H. (1991). Branching morphogenesis of mouse salivary epithelium in basement membrane-like substratum separated from mesenchyme by the membrane filter. Development 111, 327–335.

Wang, S., Matsumoto, K., Lish, S.R., Cartagena-Rivera, A.X., and Yamada, K.M. (2021). Budding epithelial morphogenesis driven by cell-matrix versus cell-cell adhesion. Cell 184, 3702–3716 e3730.

Wang, S., Sekiguchi, R., Daley, W.P., and Yamada, K.M. (2017). Patterned cell and matrix dynamics in branching morphogenesis. J Cell Biol 216, 559–570.

Wei, C., Larsen, M., Hoffman, M.P., and Yamada, K.M. (2007). Self-organization and branching morphogenesis of primary salivary epithelial cells. Tissue Eng 13, 721–735.

Williams, E.J., Furness, J., Walsh, F.S., and Doherty, P. (1994). Activation of the FGF receptor underlies neurite outgrowth stimulated by L1, N-CAM, and N-cadherin. Neuron 13, 583–594.

Yayon, A., Klagsbrun, M., Esko, J.D., Leder, P., and Ornitz, D.M. (1991). Cell surface, heparin-like molecules are required for binding of basic fibroblast growth factor to its high affinity receptor. Cell 64, 841–848.

Yurchenco, P.D., and Patton, B.L. (2009). Developmental and pathogenic mechanisms of basement membrane assembly. Curr Pharm Des 15, 1277–1294.

